# 3-Methylpentanoic acid from *Bacillus safensis* suppresses wheat blast disease by targeting UDP-glucose 4-epimerase

**DOI:** 10.64898/2026.02.23.707562

**Authors:** Md. Shahrear Parvaj Sujon, Soharth Hasnat, Rojana Binte Azad, Dipali Rani Gupta, Md Tofazzal Islam

## Abstract

Wheat blast, caused by *Magnaporthe oryzae Triticum* (MoT), is a devastating disease threatening global food security. Current reliance on chemical fungicides is unreliable due to the development of resistant MoT populations, highlighting the need for safe and eco-friendly alternatives. Naturally occurring volatile organic compounds (VOCs) possess potent antifungal potential thereby inhibiting phytopathogen. In this study, we investigated the fungicidal potential of 3-methylpentanoic acid (3-MP), a VOC produced by *Bacillus safensis* and also found in snake-fruit aroma, on MoT pathogen. In vitro assays revealed dose–dependent inhibition of MoT mycelial growth, conidiogenesis, conidial germination, and appressorium formation, with complete suppression achieved at 100–125 µM. Detached leaf, seedling, and spike assays demonstrated robust preventive and curative protection, highlighting translational potential under field conditions. Mechanistic investigations showed that 3–MP compromises membrane integrity, as confirmed by fluorescein diacetate staining, and targets UDP–glucose 4–epimerase (UGE), a key enzyme required for galactose metabolism and cell wall integrity in fungi. Molecular dynamics simulations revealed stable binding of 3–MP within the NAD⁺–associated Rossmann fold of UGE, sterically blocking substrate access and perturbing NAD⁺ orientation. RT-PCR gene expression analysis corroborated this model, showing early induction followed by repression of UGE expression, consistent with collapse of UDP–glucose metabolism and impaired cell wall biosynthesis. This study for the first time identified 3-MP as a natural inhibitor of UGE and provided new insight into the antifungal mechanism of the compound, highlighting its potential for integrated wheat blast management.

**Importance:** Wheat blast, caused by *Magnaporthe oryzae Triticum* (MoT), poses a catastrophic threat to global food security, particularly in South America, Africa, and South Asia. With MoT rapidly developing resistance to conventional chemical fungicides, there is an urgent need for sustainable, eco-friendly alternatives. Our study identifies 3-methylpentanoic acid (3-MP), a volatile organic compound produced by *Bacillus safensis*, as a potent antifungal agent against MoT. We demonstrate that 3-MP inhibits multiple life stages of the pathogen, including mycelial growth, conidiation, and appressorium formation. Furthermore, we provide molecular insights through molecular docking and MD simulations, identifying UDP-glucose 4-epimerase as a likely target of 3-MP. By revealing a dual-action (preventive and curative) natural compound and its potential mechanism, this work offers a promising blueprint for developing bio-based fungicides to combat devastating plant diseases while reducing the environmental footprint of agriculture.

## Introduction

Wheat blast, caused by the filamentous fungus *Magnaporthe oryzae Triticum* (MoT), represents one of the most destructive threats to global wheat production and food security. Since its first emergence in Brazil in 1985, the disease has rapidly expanded across South America and subsequently into Asia and Africa, including Bangladesh, highlighting its transcontinental dissemination and epidemic potential (1–5). Projections suggest that as temperatures and relative humidity increase globally, the disease could spread from tropical regions to Australia and other South American countries, potentially causing a 13% loss in global wheat yield (6). MoT spreads efficiently through infected seeds and airborne conidia, infecting all above-ground plant parts and frequently causing catastrophic yield losses under favorable environmental conditions (2, 7, 8).

Current disease management relies heavily on synthetic fungicides; however, their prolonged use has raised serious concerns regarding the development of pathogen resistance, environmental persistence, and adverse effects on human health (9–12). While breeding for resistant cultivars is a goal, it remains constrained by limited genetic sources, long development timelines, and the complex hexaploid nature of wheat, which complicates genome editing strategies (13–15). Furthermore, established resistance often breaks down at higher temperatures. These multifaceted challenges necessitate the development of alternative, environmentally compatible antifungal approaches.

Volatile organic compounds (VOCs) produced by microorganisms and plants have emerged as promising antifungal agents due to their low molecular weight (<300 Da), biodegradability, and superior ability to diffuse through air and soil matrices (16–18). Acting as infochemicals, VOCs can suppress pathogen growth, interfere with development, and induce systemic resistance in plants without leaving toxic residues. Among these, small branched-chain fatty acids have gained attention for their potent antifungal properties (19, 20). Notably, we recently identified 3-MP in the volatile profiles of *Bacillus safensis,* which inhibits phytopathogenic fungi, *Pyricularia oryzae* and *Botrytis cinerea* (21, 22). However, underlying molecular mechanism of inhibitory activity against fungi are poorly understood. Beyond its new antimicrobial activity, 3-MP naturally occurs in edible plants, such as snake fruit (23), and is classified as “Generally Recognized as Safe” (GRAS) by the U.S. Food and Drug Administration (24), highlighting its suitability for agricultural application.

Targeting essential fungal enzymes or virulence-associated proteins represents a rational strategy for developing next-generation antifungal agents with high specificity. UDP–glucose 4–epimerase (UGE) is a key metabolic enzyme that catalyzes the reversible interconversion of UDP-glucose and UDP-galactose, a precursor required for galactose metabolism and fungal cell wall biosynthesis. Because UDP-galactose contributes to structural polysaccharides critical for hyphal stability, UGE activity is indispensable for maintaining cell wall integrity (CWI). Inhibition of UGE disrupts carbohydrate flux and weakens cell wall assembly, making it an attractive antifungal target (25, 26). Similarly, fungal chitinases, transketolase (TKL), and catalase-peroxidase (KatG2) are essential for growth, pathogenicity, and counteracting host-generated oxidative stress (27–29).

Although we recently reported the antifungal activity of 3-MP (21, 22), its specific efficacy against the wheat blast pathogen MoT and the underlying molecular mechanisms of its action remains to be elucidated. In the current study, we integrated in vitro phenotypic and molecular assays with computational molecular docking and dynamics simulations to investigate the interaction of 3-MP with key MoT proteins involved in cell wall biosynthesis and pathogenicity. The primary objective of this study was to evaluate the antifungal efficacy and biocontrol potential of 3-MP against the wheat blast pathogen, while elucidating its molecular mode of action through integrated *in silico* predictions and transcriptional validation via real-time PCR. Our results demonstrate that 3-MP suppresses wheat blast disease *in planta* by preferentially targeting UGE, which leads to the repression of its expression, impairment of cell wall biosynthesis, and subsequent fungal cell death. This study provides novel mechanistic insights into the potential of 3-MP as a sustainable biorational natural products potential to control fearsome wheat blast disease.

## Results

### 3-MP inhibited mycelia growth of MoT

To evaluate the inhibitory effect of 3-MP on the vegetative growth of MoT, dual-plate assays were performed. The results demonstrated that 3-MP suppresses mycelial growth in a dose-dependent manner. Complete inhibition (100 ± 0%) was achieved at 125 µM, comparable to the synthetic fungicide Nativo® 75 WG (**Fig. 1A**). At 7 days of incubation, the highest inhibition was observed at 125 µM (100±0%), followed by 100 µM (70.07±1.16%), 75 µM (63.00±1.20%), and 50 µM (33.6±1.26%), indicating a positive correlation between 3-MP concentration and fungal growth suppression (**Fig. 1B**). After 14 days, following removal of 3-MP at day 7, complete inhibition persisted at 125 µM (**Fig. 1C**). In contrast, inhibition at 100 µM declined to 40.8 ± 0.78%, while 75 µM and 50 µM showed only 10.2 ± 0.49% and 5.29 ± 0.59% suppression, respectively (**Fig. 1D**). These results demonstrate that 3-MP is fungicidal and effective at lower (125 µM) concentrations. We have also tested this compound against one of our Bangladeshi rice blast isolate RbMe1819-3. Our result demonstrated that 3-MP completely suppresses mycelial growth of rice blast fungus at 125 µM concentration (**Fig. S1**).

**Figure 1.**
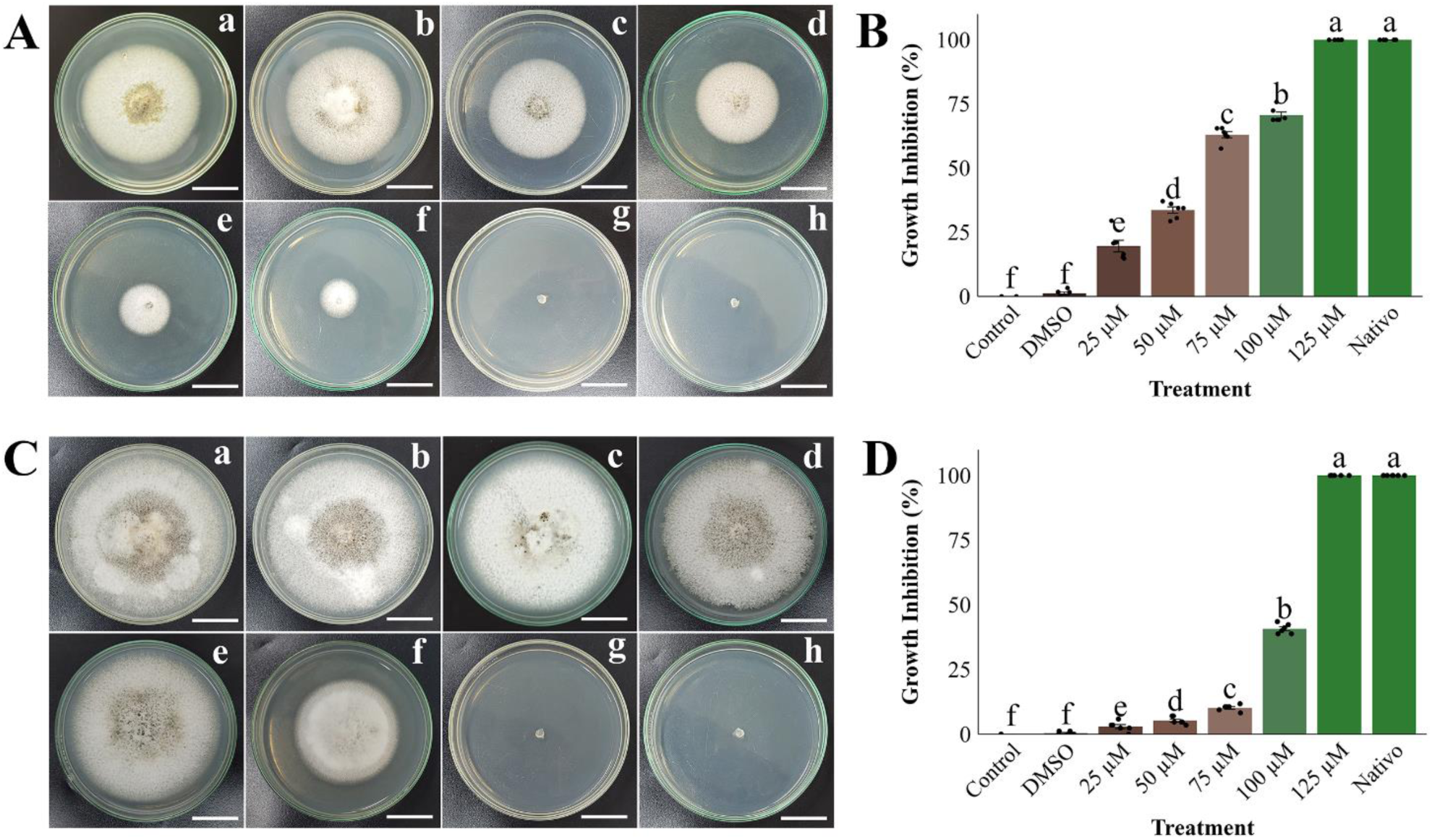
Inhibitory effects of 3-MP on mycelial growth of MoT. **(A)** Micrograph representing the suppression of mycelial growth of 3-MP in PDA; **(B)** MoT mycelial growth inhibition (%) by 3-MP over untreated control. **(C)** Fungicidal inhibitory effects of 3-MP on mycelial growth of MoT after volatile containing filter paper removed **(D)** MoT mycelial growth inhibition (%) by 3-MP over untreated control after removing 3-MP containing filter paper disc. (a) Control, (b) DMSO 1%, (c) 25 µM, (d) 50 µM, (e) 75 µM, (f) 100 µM, (g) 125 µM, (h) Nativo® WG75. Data were recorded 7 days (**A** and **B**) and 14 days (**C** and **D**) after treated and incubated at 25(±2) °C. Error bars represent the standard error calculated from six biological replicates. Different letters above the bars indicate statistically significant differences between treatments (P ≤ 0.01), as determined by one–way ANOVA followed by the Tukey–HSD post hoc test. Bar = 2 cm

### Conidiogenesis is inhibited by 3-MP

Conidiogenesis is a critical phase in the MoT infection cycle (14). To assess the efficacy of 3-MP in disrupting this process, we compared its inhibitory effects against the commercial fungicide Nativo® 75 WG. Both 3-MP and Nativo® 75 WG significantly suppressed conidial production relative to the untreated controls. Inhibition by 3-MP was concentration-dependent, with progressive suppression at 50, 100, and 150 µM (**Fig. 2A**). At 150 µM, MoT produced only 1.66 ± 0.33 conidia/mm², and microscopic examination revealed broken hyphal tips and complete absence of conidiophores. In contrast, the control plate treated with sterile water containing 1% DMSO yielded ∼1922.66 ± 16.33 conidia/mm² (**Fig. 2B**).

**Figure 2.**
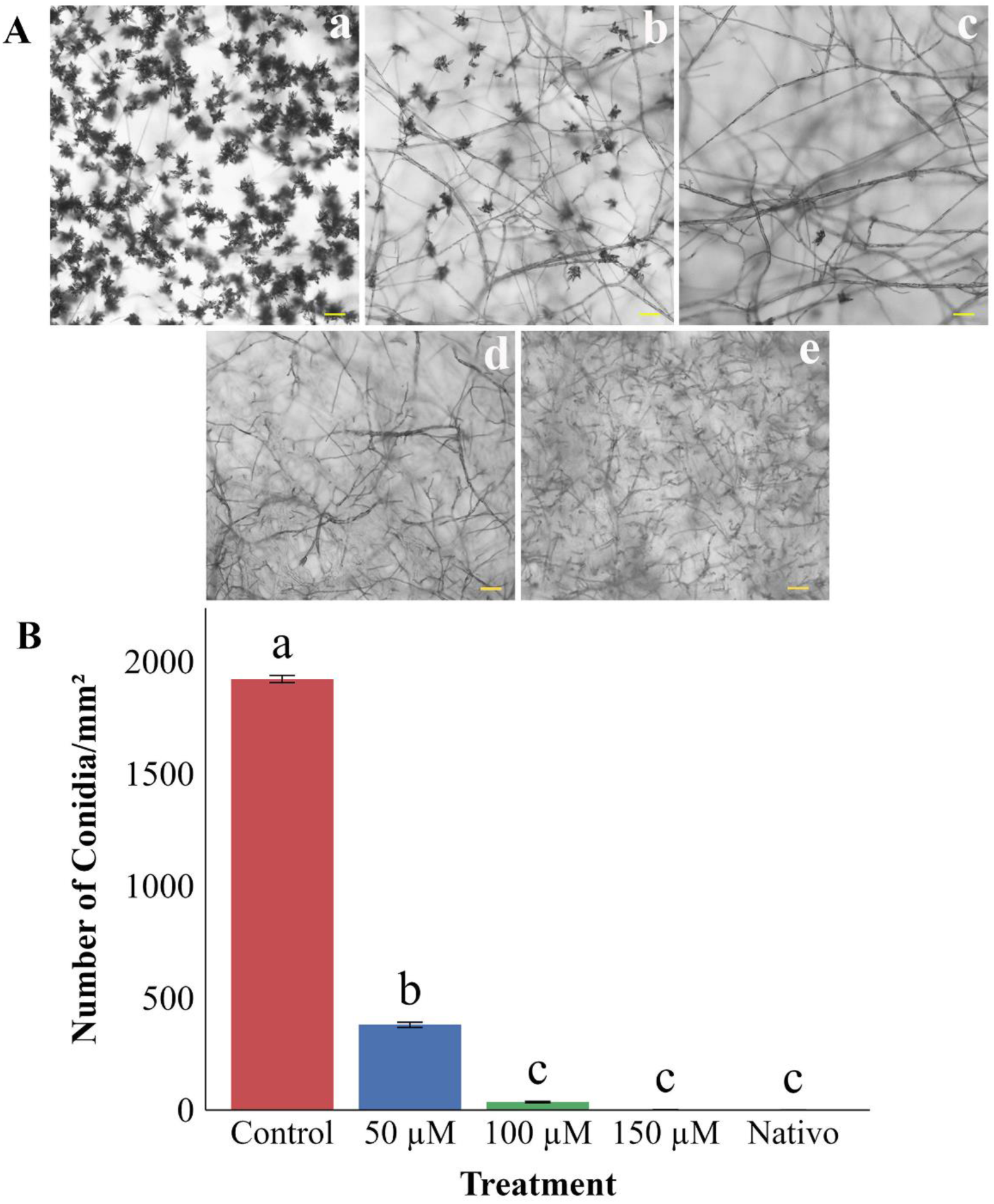
Effects of 3-MP and the commercial fungicide Nativo® 75 WG on the suppression of conidiogenesis of MoT. **(A)** Microscopic visualization of conidial development under different concentrations. (a) Control, (b) 50 µM, (c) 100 µM, (d) 150 µM, (f) Nativo® 75 WG. **(B)** Quantification of conidia density (number of conidia/mm²) across treatments. Error bars represent mean ± SE. Treatments sharing the same letter are not significantly different (p < 0.01), based on Tukey’s HSD test following one-way ANOVA. Bar = 50 µM

### 3-MP suppresses conidia germination and appressorium formation

The airborne conidia of MoT are the primary agents for dispersal and host infection. Because conidial germination is the first critical step in initiating the infection process (2, 14). We evaluated the inhibitory potential of 3-MP and our results revealed that 3-MP significantly suppresses both conidial germination and appressorium formation in a concentration-dependent manner. Conidia demonstrated greater sensitivity to 3-MP compared to mycelia, with germination monitored at 3, 6, 9, 12, and 24 hours (h) post-incubation. In water control, conidia achieved 100% germination, with normal germ tube development at 9-24 h of incubation at 25°C in the dark (**Fig. 3A** and **Table 1**). In contrast, 3-MP markedly reduced germination and disrupted development. At 9 h, germination declined to 80 ± 2.31%, 13.7 ± 1.67%, and 0 ± 0% at 25, 50, and 100 µM, respectively (**Fig. 3B** and **Table 1**). Treated conidia exhibited short germ tubes and abnormal transitions, with cell wall rupture evident at 100 µM. Developmental defects persisted at 12 h, and by 24 h, germination was reduced to 1.67 ± 0.33%, with complete inhibition of appressorium formation.

**Figure 3.**
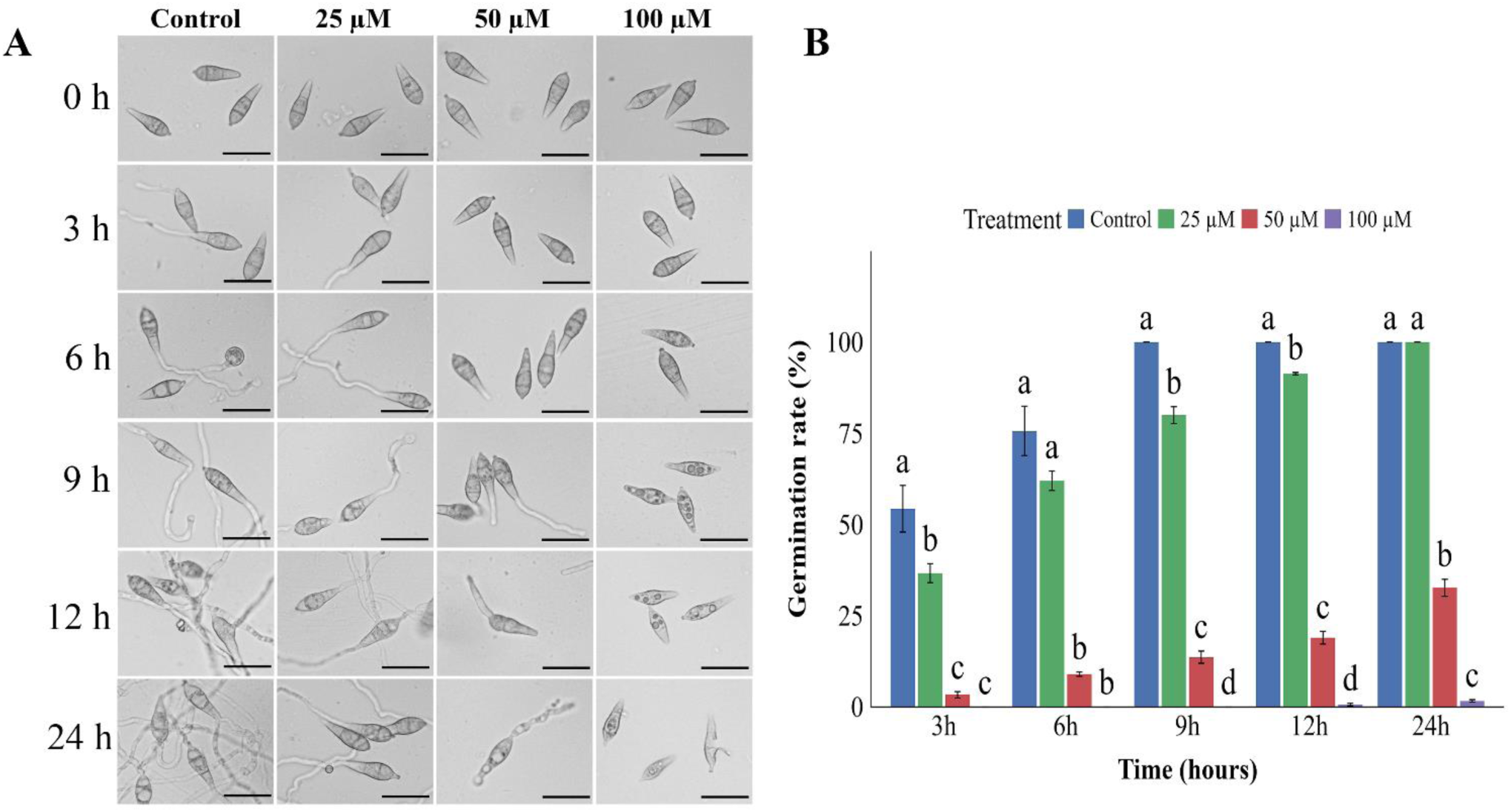
Effect of 3-MP on spore germination and morphology alteration **(A)** Time-course microscopic observations of MoT conidia exposed to 3-MP at 25 µM, 50 µM, and 100 µM concentrations, compared to untreated control. Images were captured at 0, 3, 6, 9, 12, and 24 hours post-incubation. **(B)** Quantitative analysis of conidial germination rate (%) under different treatments over time. Each data point represents the mean ± SE from three independent replicates, with 100 conidia counted per replicate. Different letters above bars indicate statistically significant differences among treatments at each time point (Tukey’s HSD, p < 0.01). Bar = 25 µm.

**Table 1.** Effects of 3-MP on germination of conidia and morphology of germ tubes and appressoria of MoT at 25 µM, 50 µM, and 100 µM *in vitro*.

| Compound | Time (h) | Germinated (% ± SE) | Major morphological changes occurred in the treated conidia |
| --- | --- | --- | --- |
| Control | 0 | $0 \pm 0^a$ | No germination |
| | 3 | $54.33 \pm 6.38^a$ | Normal germ tube initiation |
| | 6 | $75.7 \pm 6.76^a$ | Germinated with normal germ tube and normal appressoria developed |
| | 9 | $100 \pm 0^a$ | Extended hyphal growth |
| | 12 | $100 \pm 0^a$ | Significant mycelial elongation |
| | 24 | $100 \pm 0^a$ | Fully developed hyphal networks |
| 25 μM | 0 | $0 \pm 0^a$ | No germination |
| | 3 | $36.67 \pm 2.60^b$ | Germinated with normal germ tube |
| | 6 | $62 \pm 2.64^a$ | No appressoria formed |
| | 9 | $80 \pm 2.31^b$ | Hyphal growth was continued but branching was limited |
| | 12 | $91.33 \pm 0.33^b$ | Hyphal growth was reduced with fewer branches |
| | 24 | $100 \pm 0^a$ | Developed hyphal networks but appressoria formed |
| 50 μM | 0 | $0 \pm 0^a$ | No germination |
| | 3 | $3.33 \pm 0.88^c$ | No germination observed |
| | 6 | $9 \pm 0.57^b$ | Minor germ tube initiation |
| | 9 | $13.67 \pm 1.67^c$ | Germinated conidia had short germ tube |
| | 12 | $19 \pm 1.73^c$ | Conidia with limited growth and no appressoria formed |
| | 24 | $32.67 \pm 2.33^b$ | Deformed germ tube (swollen) and no mycelial growth |
| 100 μM | 0 | $0 \pm 0^a$ | No germination |
| | 3 | $0 \pm 0^c$ | No germination |
| | 6 | $0 \pm 0^b$ | Conidia remained inactive |
| | 9 | $0 \pm 0^d$ | Zero appressoria formed; zero hyphal growth |
| | 12 | $0.67 \pm 0.33^d$ | Conidia vacuolization was observed |
| | 24 | $1.67 \pm 0.33^c$ | Conidia appeared shriveled, showing cell wall leakage. |

### 3-MP Suppresses disease lesion development on detached leaves of wheat

To determine whether 3-MP could protect wheat foliage from MoT infection, we performed artificial inoculation on detached wheat leaves using MoT conidia. Our results demonstrated that 3-MP significantly reduced blast symptoms in a concentration-dependent manner. At 25 and 50 µM, average lesion areas were 0.45 ± 0.037 cm² and 0.041 ± 0.004 cm², respectively. Lesions were absent at 100 µM (**Figs. 4, A** and **B**). In contrast, inoculated control leaves developed typical blast lesions with an average size of 0.64 ± 0.05 cm².

**Figure 4.**
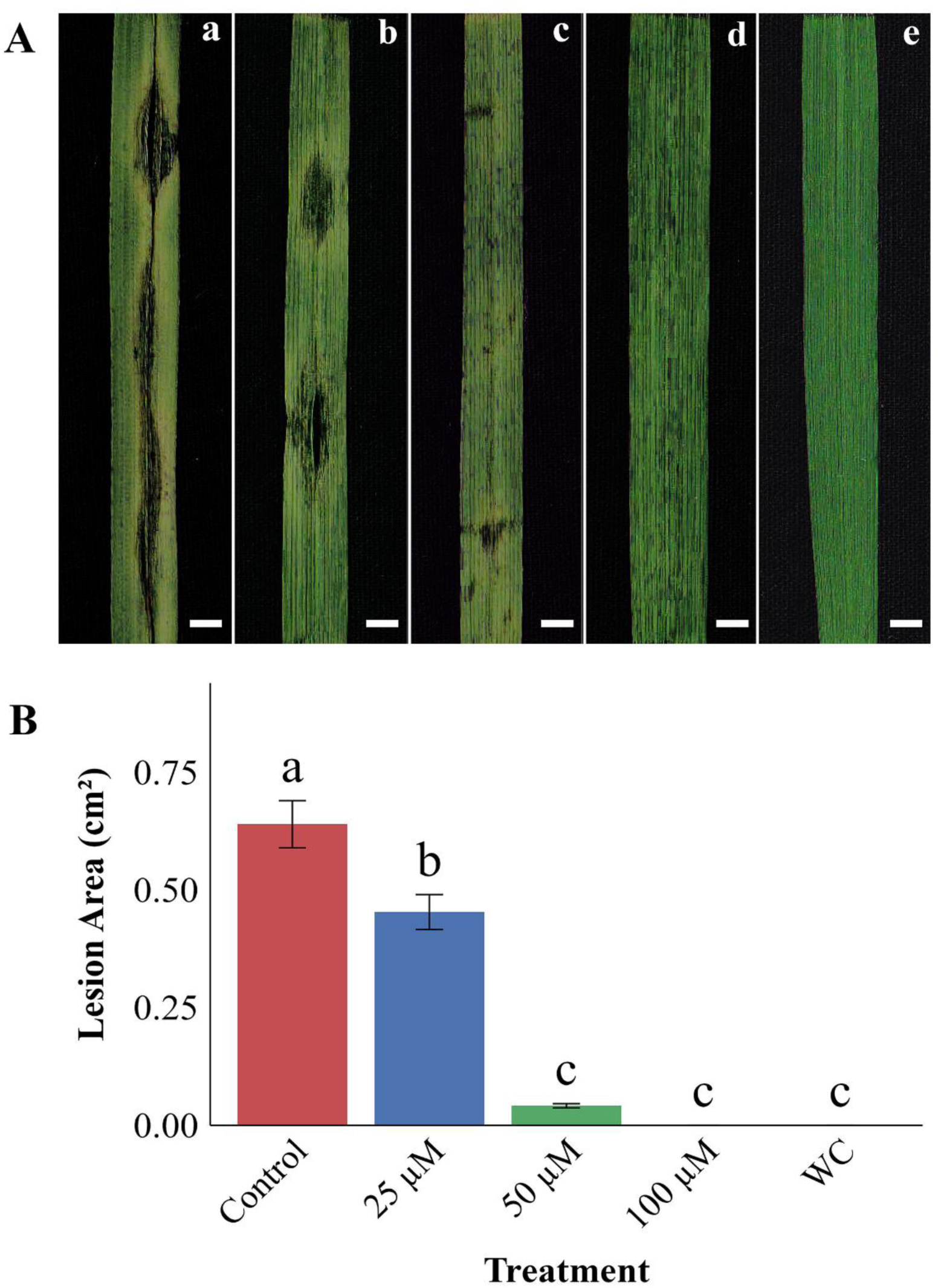
Effects of 3-MP on the suppression of lesion formation in detached wheat leaves by MoT **(A)** Development of blast lesion on 3-MP treated and untreated wheat leaves. (a) Control, (b) 25 µM, (c) 50 µM, (d) 100 µM, (e) sterile water treated **(B)** Bar graph represent the diameter of lesions (cm^2^) in different treatments. Bars represent mean ± SE from three biological replicates. Different letters above bars indicate statistically significant differences among treatments (Tukey’s HSD, p < 0.01). Bar = 5 mm

### In planta wheat blast disease suppression by 3-MP Preventive effect

To evaluate the biocontrol potential of 3-MP against wheat blast, seedlings were treated with the compound prior to artificial inoculation with MoT conidia. Preventive application of 3-MP 48 h before inoculation significantly reduced disease symptoms in seedlings challenged with the virulent strain BTJP 4 (5). At 32 µM, disease incidence and severity were 68.4 ± 2.17% and 39.2 ± 3.09%, respectively, whereas at 64 µM they declined to 27.2 ± 3.14% and 11.7 ± 1.88% (**Figs. 5, A** and **B**). Untreated controls exhibited the highest disease incidence (98.1 ± 1.85%) and severity (63.9 ± 2.62%), while healthy controls showed no symptoms (**Table 2**).

**Figure 5.**
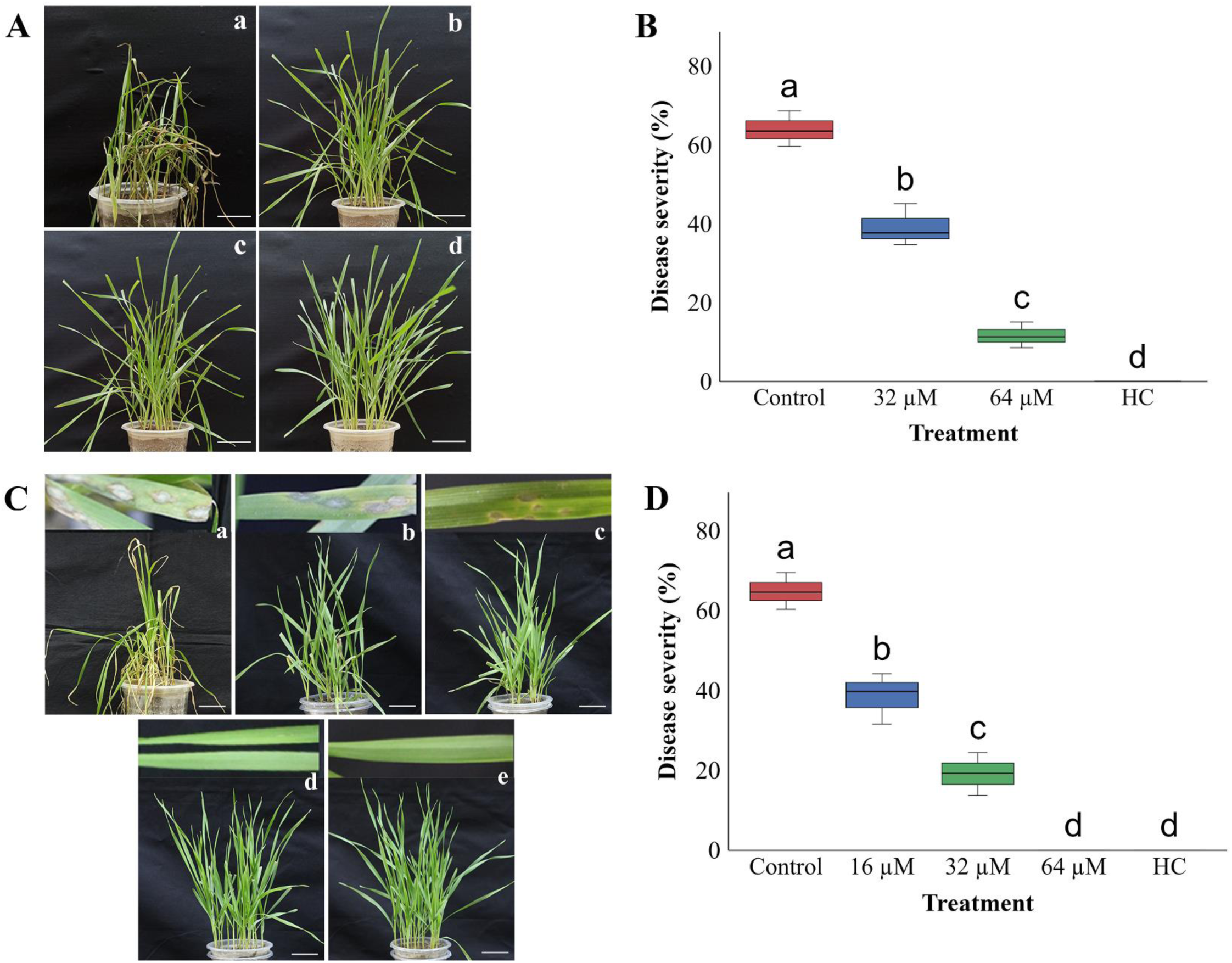
Preventive and curative effects of 3-MP on blast disease in wheat seedlings. **(A)** Effect of 3-MP on controlling seedling leaf blast caused by MoT (preventive assay). (a) Control, (b) 32 µM, (c) 64 µM, (d) healthy control. **(B)** Blast disease severity (%) in seedlings pre-treated with 3-MP, relative to the untreated control. **(C)** Effect of 3-MP on controlling seedling leaf blast caused by MoT (curative assay). (a) Control, (b) 16 µM, (c) 32 µM, (d) 64 µM, (e) healthy control. **(D)** Blast disease severity (%) in post-inoculation treatments compared to untreated control. Error bars represent mean ± SE from three biological replicates. Different letters indicate statistically significant differences among treatments (Tukey’s HSD, p < 0.01). Bar = 3.5 cm

**Table 2.** Effect of 3-MP in suppression of wheat blast disease development in artificially inoculated wheat seedlings. Means ± standard errors having a common letter are not significantly different at the 0.1% level of significance.

| Parameter | Healthy<br>Control | Preventive |  |  | Curative |  |  |  |
| --- | --- | --- | --- | --- | --- | --- | --- | --- |
| | | Untreated<br>Control | 32 μM | 64 μM | Untreated<br>Control | 16 μM | 32 μM | 64 μM |
| Disease<br>incidence<br>(%) | 0±0.00<sup>d</sup> | 98.1±1.85<sup>a</sup> | 68.4±2.17<sup>b</sup> | 27.2±3.14<sup>c</sup> | 98±1.96<sup>a</sup> | 76.5±3.02<sup>b</sup> | 53.7±4.07<sup>c</sup> | 0±0.00<sup>d</sup> |
| Disease<br>Severity<br>(%) | 0±0.00<sup>d</sup> | 63.9±2.62<sup>a</sup> | 39.2±3.09<sup>b</sup> | 11.7±1.88<sup>c</sup> | 64.8±2.66<sup>a</sup> | 38.5±3.69<sup>b</sup> | 19.1±3.10<sup>c</sup> | 0±0.00<sup>d</sup> |

### Curative effect

To assess the curative efficacy of 3-MP, the compound was applied 48 h post-inoculation of wheat seedlings with MoT conidia. At doses of 16 and 32 µM, disease incidence was 76.5 ± 3.02% and 53.7 ± 4.07%, respectively, with corresponding severities of 38.5 ± 3.69% and 19.1 ± 3.10% (**Figs. 5, C** and **D**). Notably, complete suppression of the disease was achieved at 64 µM, with no visible blast symptoms observed (**Table 2**).

### Application of 3-MP effectively suppresses wheat blast disease development in wheat head

Wheat blast is primarily a head-infecting disease, with the flowering stage (anthesis) representing the most vulnerable phase of crop development. To evaluate the practical application of 3-MP as a bio-fungicide, we tested its efficacy on wheat spikes during anthesis. 3-MP provided robust protection; for instance, treatment with 16 µM reduced disease incidence to just 6.0%, compared with 86.9% in untreated controls. Disease severity was similarly suppressed in a dose-dependent manner; plants treated with 8 and 16 µM exhibited severities of 29.9 ± 3.62% and 1.5 ± 0.84%, respectively, whereas untreated plants reached 75.3 ± 3.15% (**Figs. 6A**, **B** and **Table 3**).

**Figure 6.**
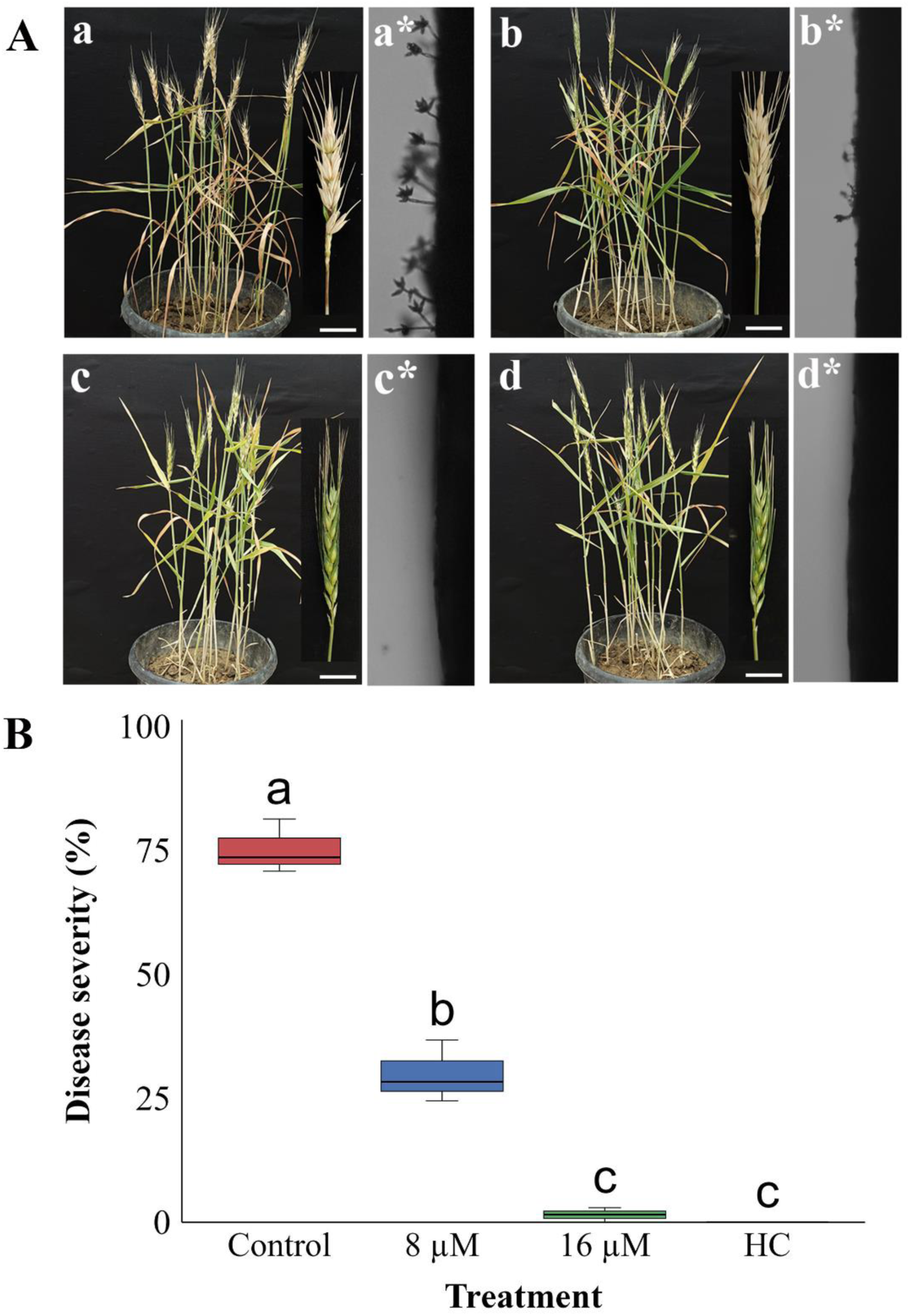
Effect of 3-MP on wheat head blast suppression. **(A)** Photographs showing the disease severity in wheat heads following MoT inoculation and 3–MP treatment (a) Control, (a*) microscopic image of control spike (b) 8 µM, (b*) microscopic image of 8 µM 3-MP treated spike (c) 16 µM, (c*) microscopic image of 16 µM 3-MP treated spike (d) healthy control (d*) microscopic image of control spike and **(B)** Wheat head blast disease severity (%) under different treatment conditions. Error bars represent mean ± SE from three biological replicates. Different letters indicate statistically significant differences among treatments (Tukey’s HSD, p < 0.01). Bar = 4.5 cm

**Table 3.** Effect of 3-MP in suppression of wheat blast disease development in artificially inoculated wheat spike in green house condition. Means ± standard errors having a common letter are not significantly different at the 0.1% level of significance

| Parameter | Healthy Control | Untreated Control | 8 μM | 16 μM |
| --- | --- | --- | --- | --- |
| % Disease incidence | 0± 0.00<sup>c</sup> | 86.9 ± 4.82<sup>a</sup> | 48.0± 2.67<sup>b</sup> | 6.0 ± 3.40<sup>c</sup> |
| % Disease Severity | 0± 0.00<sup>c</sup> | 75.3 ± 3.15<sup>a</sup> | 29.9± 3.62<sup>b</sup> | 1.5 ± 0.84<sup>c</sup> |

### 3-MP disrupted membrane integrity and compromised the viability of MoT cells

To elucidate the mechanism underlying hyphal growth inhibition, we employed fluorescein diacetate (FDA) staining to assess cellular viability and membrane integrity. Our results revealed distinct differences between untreated and 3-MP–treated MoT hyphae. Untreated hyphae exhibited robust green fluorescence, indicative of metabolic activity and intact cytoplasmic membranes. Conversely, treatment with 3-MP at 50–100 µM resulted in a progressive decline in fluorescence intensity, signifying compromised membrane permeability and diminished viability (**Fig. 7**). At the 125 µM threshold, fluorescence was entirely abolished, reflecting irreversible membrane damage and complete loss of hyphal viability.

**Figure 7.**
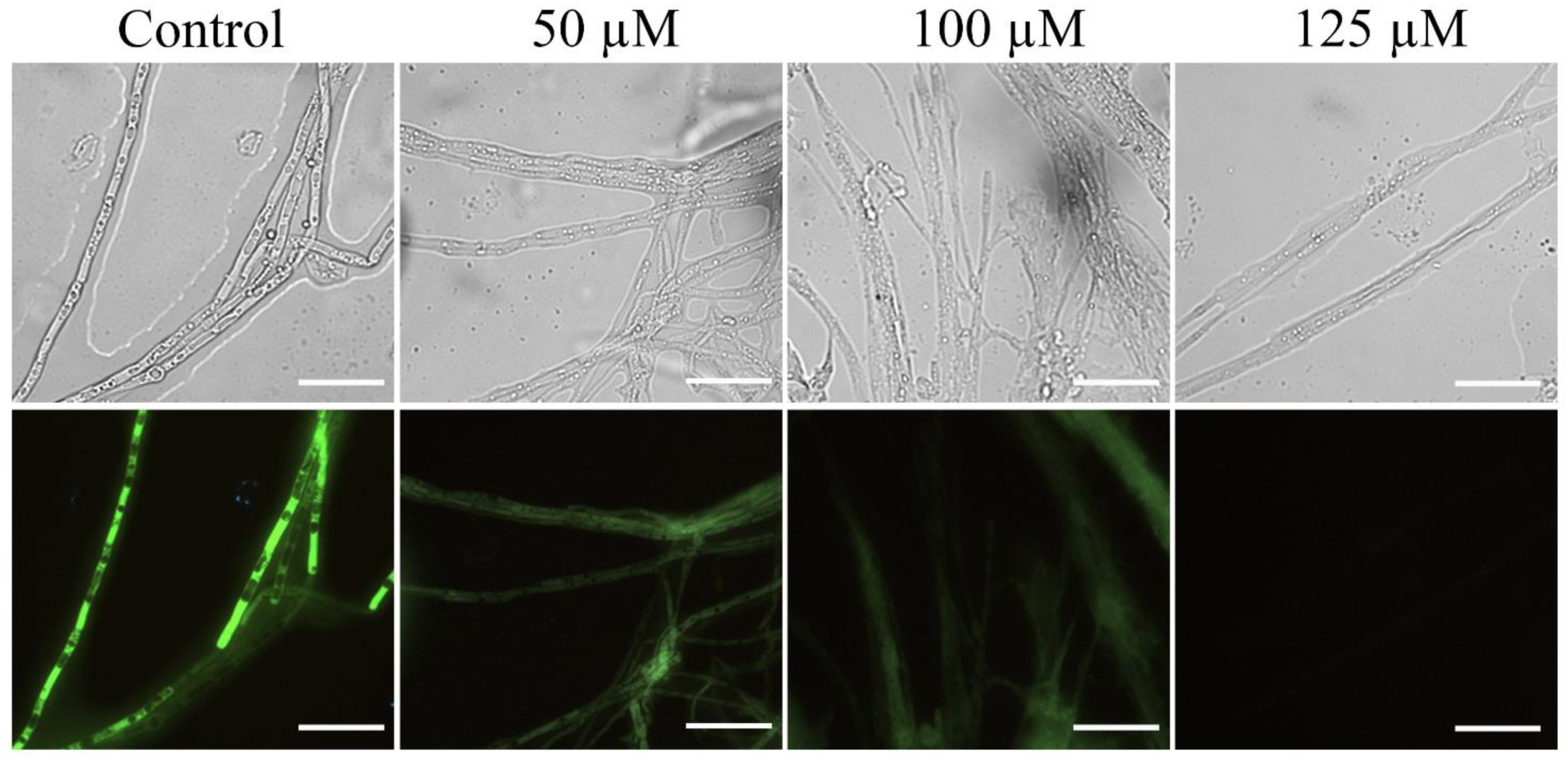
Fluorescent detection of MoT viability after 3-MP treatment using FDA staining. Intact, viable hyphae emitted strong green fluorescence, while weakened signals in treated samples highlighted 3-MP–mediated inhibition. Images were obtained at 40X with a Zeiss Axiocam ERc 5s camera. Bar = 20 µm

### In silico analysis reveals stable interactions between 3-MP and essential MoT proteins

To identify the molecular targets of 3-MP among essential enzymes involved in MoT membrane and cell wall biosynthesis, we employed a high-throughput computational screening approach. Screening of 196 essential MoT proteins identified four primary targets: catalase-peroxidase (MagKatG2, PDB ID: 3UT2), UDP-glucose 4-epimerase (UGE, UniProt ID: G4MX57), chitin synthase 1 (Chs1, UniProt ID: G4MVT6), and transketolase (TKL, UniProt ID: G4MRY4). These candidates were prioritized based on their high binding affinities and robust interaction profiles with 3-MP (**Fig. 8** and **Fig. S2**). To evaluate the thermodynamic stability of these protein-ligand complexes, we performed molecular dynamics simulations (MDS). The UGE–3-MP complex displayed the highest structural stability, maintaining Root Mean Square Deviation (RMSD) fluctuations between 1–2.5 Å, which indicates a highly favorable and rigid binding conformation. While MagKatG2–3-MP showed moderate stability (RMSD: 1–4 Å), the Chs1 and TKL complexes exhibited broader fluctuations (RMSD: 1–5 Å and 1–6 Å, respectively) (**Fig. 8** and **Fig. S2**). Collectively, these RMSD profiles identify UGE as the most probable high-affinity target, though the relative stability of all four complexes suggests a potential multi-target mechanism contributing to the antifungal activity of 3-MP.

**Figure 8.**
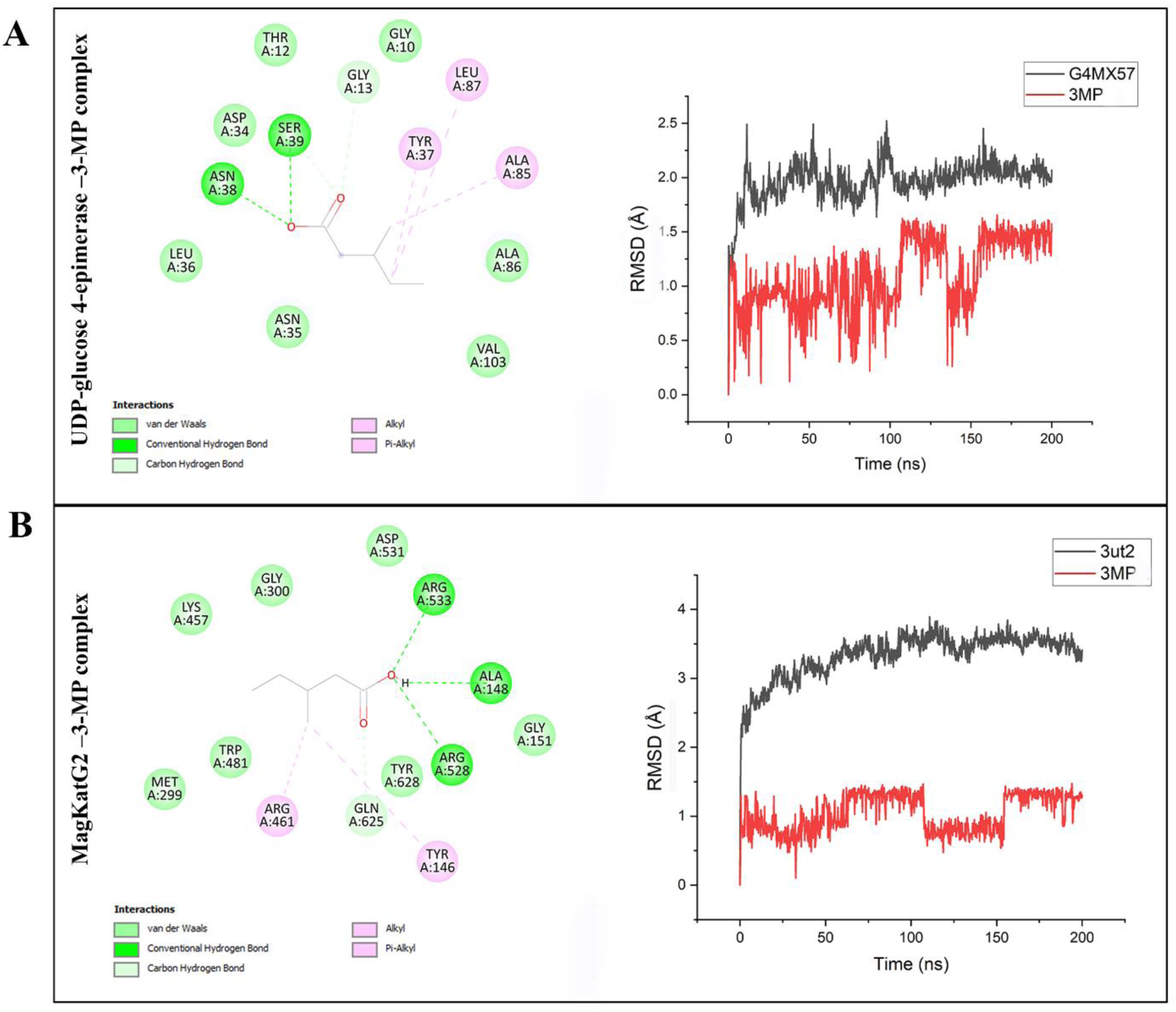
Molecular interaction and stability analysis of 3-MP with key MoT proteins. **(A)** Interaction profile of 3-MP with UDP-glucose 4-epimerase, showing hydrogen bonding, van der Waals, and alkyl interactions with active site residues. The RMSD plot indicates stable binding of 3-MP to the protein over the simulation period. **(B)** Interaction profile of 3-MP with MagKatG2, highlighting key residue contacts and binding forces. RMSD analysis reveals enhanced structural stability of the 3-MP–protein over the simulation period.

### 3-MP represses *UGE* expression involved in UDP-sugar metabolism and cell wall biosynthesis in MoT

To validate the *in silico* predictions indicating that 3-MP binds to UGE, we characterized the *in vivo* transcriptional response of *UGE* and other key genes MoT using RT-qPCR. Fungal cultures were exposed to sublethal (100 µM) and lethal (150 µM) concentrations of 3-MP, and the expression profiles of four essential genes *UGE*, *Chs1*, *MagKatG2*, and *Transketolase* were monitored at 24 and 48 h post-exposure. The expression of *UGE* exhibited a distinct temporal shift. At 24 h, the gene was moderately upregulated, with Log_2_ fold changes of approximately 0.95 and 0.78 under 100 µM and 150 µM treatments, respectively. However, by 48 h, *UGE* expression was significantly repressed at both concentrations, showing negative Log_2_ fold changes of −0.44 (100 µM) and −0.58 (150 µM) (**Fig. 9A**). This biphasic pattern initial induction followed by sustained repression corroborates our molecular dynamics model and is consistent with a collapse in UDP-glucose metabolism. In contrast, *Chs1* expression was consistently upregulated at both time points. At 24 h, transcript levels increased substantially, reaching Log_2_ fold changes of 3.4 (100 µM) and 3.8 (150 µM). Although the magnitude of induction slightly decreased by 48 h (2.9 at 100 µM; 3.4 at 150 µM), expression remained elevated, likely representing a compensatory response to cell wall stress (**Fig. 9B**). Distinct temporal patterns were also observed for *MagKatG2* and *Transketolase*. *MagKatG2* showed minimal changes at 24 h (Log2 fold changes of ∼0.9–1.1) but exhibited a marked induction at 48 h, peaking at 3.5 (100 µM) and 2.9 (150 µM). Similarly, *Transketolase* expression varied by dose at 24 h, with Log2 fold changes of 2.6 (100 µM) and 0.7 (150 µM). By 48 h, however, *Transketolase* was strongly upregulated regardless of concentration, reaching Log2 fold changes of 4.7 and 4.9 for the sublethal and lethal doses, respectively (**Figs. 9C** and **D**). These significant increases in the expression of antioxidant and metabolic genes suggest an intensive but ultimately insufficient cellular attempt to counteract 3-MP-induced oxidative stress and metabolic imbalance.

**Figure 9.**
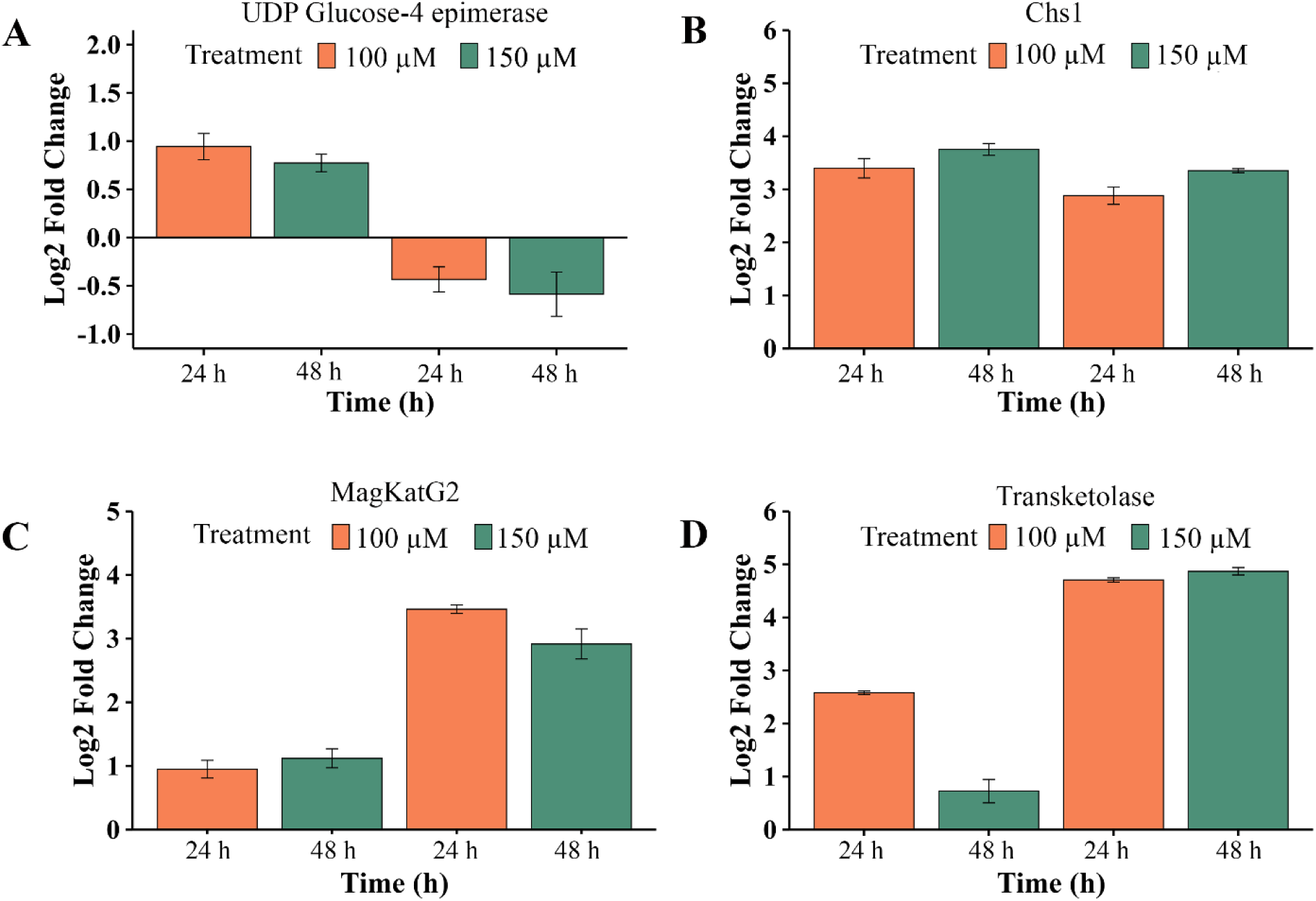
Relative expression (Log₂ fold change) of four genes in MoT following treatment with 3-MP. qRT-PCR analysis was performed for **(A)** UGE, **(B)** Chs1, **(C)** MagKatG2, and **(D)** Transketolase under sublethal (100 µM) and lethal (150 µM) concentrations at 24 h and 48 h

## Discussion

The rapid emergence of fungicide-resistant MoT populations presents a significant challenge to global wheat security, underscoring the urgent need for sustainable control strategies that transcend conventional chemistries (12). Although, recently, we reported that 3-MP is an antifungal volatile compound (21, 22), its effect on highly destructive and emerging wheat blast fungus, MoT and the precise molecular mode of action against filamentous phytopathogens has remained undefined. In this study, we integrated *in vitro* phenotypic assays with computational molecular docking, molecular dynamics (MD) simulations, and transcriptional gene expression analysis to elucidate the antifungal mechanism of 3-MP against MoT. Our findings demonstrated that 3-MP is a potent bio-based fungicide that exerts lethal effects to suppress wheat blast disease in planta by simultaneously compromising membrane integrity and disrupting essential carbohydrate metabolism likely through targeting of UGE. The discovery of this novel mode of action for a GRAS-certified flavoring agent against MoT provides a robust framework for the design of biorational fungicides. However, extensive field-scale evaluations and the development of optimized formulations are essential prerequisites to validate its efficacy and stability before 3-MP can be recommended for large-scale agronomic application to combat wheat blast. Some natural products such as VOCs (hexanoic acid, 2-methylbutanoic acid, and phenylethyl alcohol) from *Bacillus* species, antimycin A, oligomycins, lipopeptides and staurosporine have been reported as inhibitors of wheat blast fungus, however, mode of actions of these compounds are unknown (30, 31, 35, 46).

Phenotypically, 3-MP exhibited potent, dose-dependent inhibition of MoT mycelial growth, with complete suppression achieved at 125 µM (**Figs. 1, A** and **B**). This inhibitory threshold is comparable to the efficacy of the commercial fungicide Nativo® 75 WG at recommended field rates (31), positioning 3-MP as a high-potential bio-based alternative. Beyond vegetative growth, 3-MP markedly disrupted the MoT infection cycle by impairing conidiogenesis, conidial germination, and appressorium formation (**Figs. 2 and 3**). Given that conidia are the primary inoculum and germination and appressoria development are functionally indispensable for host penetration (32–34), the impairment of formation and conidia and their subsequent germination at at micromolar concentration suggests that 3-MP serves as a powerful barrier to both pathogen dissemination and host colonization. The simultaneous inhibition of mycelial expansion, conidiogenesis, conidial germination, and appressorial differentiation is essential for the comprehensive impairment of the MoT cycle (35).

A critical observation in this study was the induction of severe morphological aberrations, including hyphal tip lysis and extensive vacuolization, signaling a catastrophic collapse of cellular homeostasis. Our FDA staining assays confirmed a dose-dependent loss of fluorescence, correlating with irreversible disruption of membrane integrity (**Fig. 7**). While small-chain volatile fatty acids are known to integrate into lipid bilayers and perturb membrane fluidity (19, 33), our study suggests that the antifungal efficacy of 3-MP is not merely a consequence of non-specific surfactant-like effects, but involves highly specific protein-ligand interactions.

The identification of UGE as a high-affinity ligand for 3-MP represents a hallmark of this work, linking VOC exposure to the targeted inhibition of sugar-nucleotide metabolism. UGE is a pivotal enzyme in the Leloir pathway, catalyzing the interconversion of UDP-glucose and UDP-galactose, which are essential precursors for fungal cell wall polysaccharides and glycoprotein biosynthesis (36–38). Our *in silico* docking and MD simulations revealed that 3-MP stably occupies the N-terminal cofactor-binding cleft of UGE, forming a robust hydrogen-bond network with Ser39 and engaging the conserved glycine-rich motif (Gly10, Thr12, and Gly13) (**Fig. 8**). This binding mode sterically occludes the NAD⁺-binding pocket. Unlike traditional competitive inhibitors, 3-MP appears to inhibit UGE via cofactor displacement, a mechanism that may offer higher specificity and reduced risk of cross-resistance with current site-specific fungicides.

Another significant finding of this study is the marked transcriptional downregulation of *UGE* in 3-MP-treated MoT cultures (**Fig. 9**). This observation provides experimental validation of our computational predictions, demonstrating that 3-MP exerts a dual inhibitory effect by targeting UGE at both the structural (enzymatic) and regulatory (transcriptional) levels (**Figs. 9** and **10**). While disruption of *UGE* in *M. oryzae Oryzae* and other fungi like *Aspergillus niger* leads to cell wall defects rather than immediate lethality (27, 39), the near-complete suppression of MoT demonstrated in the current study may suggest that 3-MP likely perturbs additional essential metabolic or NAD⁺-dependent pathways beyond UDP-galactose biosynthesis. The resulting failure of cell wall biosynthesis, coupled with the collapse of central carbon metabolism and associated pathways, likely creates a synergistic “metabolic trap” that leads to fungal death. Furthermore, the significant reduction of blast lesions in detached leaves, seedlings, and in the spikes (**Figs. 4–6**) demonstrates the translational potential of 3-MP. Consistent with this, inhibition of *MoUGE1*, a homolog of *MoT UGE*, by the selective synthetic inhibitor (lig122132) has recently been shown to attenuate blast disease severity in rice through suppression of mycelial growth, appressorium formation, and host penetration, while exerting little effect on conidiation (39). In contrast, 3-MP exhibits a broader impact on *MoT* morphology, markedly inhibiting mycelial growth, conidiogenesis, and conidial germination, thereby leading to a more pronounced reduction in blast disease severity (**Figs. 2, 3 *and* 5**). Interestingly, the near-complete protection observed in curative applications compared to preventive suggests a dual mode of action of 3-MP: direct fungicidal activity and the potential priming of host innate immunity. The high volatility of 3-MP likely facilitates rapid systemic penetration into wheat tissues, similar to the action of microbial volatiles like dimethyl sulfide, which are known to induce systemic resistance (ISR) while directly suppressing pathogens (32, 40). Unlike the *MoUGE1* mutant of the rice blast pathogen *M. oryzae*, which retains the ability to produce conidia, 3-MP treatment results in a total (100%) block of conidiogenesis in *MoT*. This discrepancy suggests that 3-MP may have secondary targets beyond the UGE enzyme in *MoT*. Consequently, further investigation is required to fully elucidate the multi-target mechanism of action of 3-MP in phytopathogenic fungi (39).

**Figure 10.**
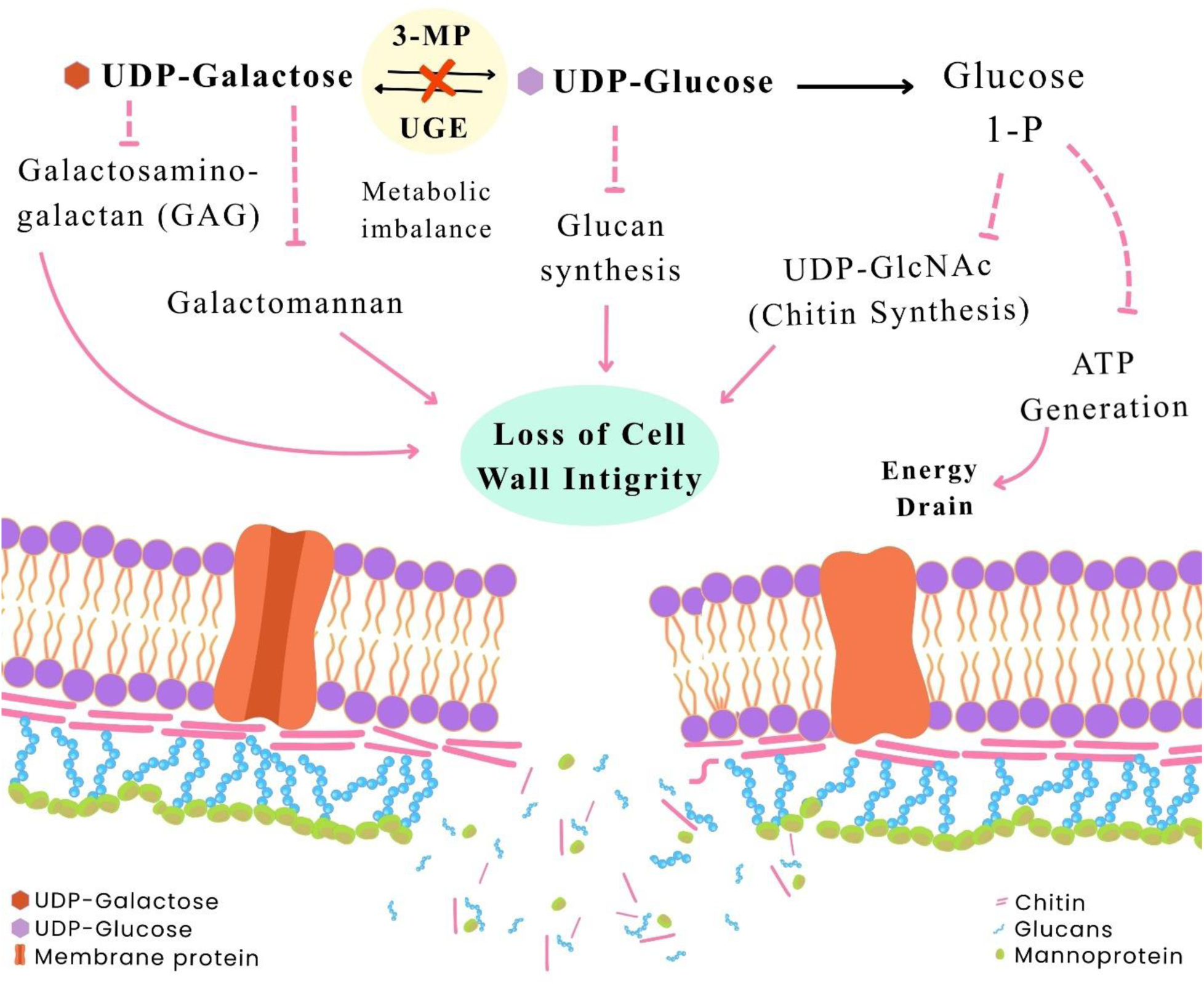
3-MP mediated inhibition of UGE impairs the Leloir pathway, disturbs nucleotide sugar balance, glucan synthesis, chitin synthesis and leads to abnormal galactomannan deposition with weakened fungal cell wall integrity

Based on *in vitro, in planta* and *in silico* study, we proposed a putative antifungal mode of action of 3-MP against MoT. Binding of UGE by 3-MP disrupts the interconversion between UDP-glucose and UDP-galactose, leading to disruption of UDP-galactose/galactose homeostasis in MoT cell. Both glucose and galactose are vital for the synthesis of glycan and galactose-containing cell wall polysaccharides. Reduced availability of any one of the sugar compromises proper cell wall assemblies, weakens structural integrity, and triggers compensatory cell wall stress responses. In parallel, perturbation of UDP-sugar homeostasis alters carbon flux from central metabolism toward nucleotide-sugar biosynthesis, creating metabolic imbalance that can impair energy utilization and biosynthetic capacity. Collectively, 3MP-mediated UGE inhibition links disruption of central carbohydrate metabolism with defective cell wall architecture, ultimately constraining fungal growth and pathogenic development (**Fig. 10**).

In summary, our results establish 3-MP as a potent bio-based fungicide for the management of the devastating wheat blast disease. Collectively, our data support a model wherein 3-MP likely to target UGE to disrupt the interconversion of essential sugar nucleotides, thereby crippling cell wall integrity and compromising metabolic flux. This mechanism highlights UGE as a highly vulnerable metabolic “Achilles’ heel” in phytopathogenic fungi. These insights position 3-MP as a promising lead compound for bio-fungicide development and provide a framework for the rational design of new agents targeting the metabolic vulnerabilities of the wheat blast pathogen. Future research utilizing targeted gene deletion and proteome-wide identification will be instrumental in further resolving the multi-layered lethal mechanism of 3-MP. Extensive field-scale evaluations are required to validate the efficacy of 3-MP under diverse environmental conditions before its recommendation as a bio-based fungicide for large-scale agronomic application.

## Materials and Methods

### Fungal Strain and Culture Conditions

The wheat blast strain BTJP 4 (5) was isolated from an infected wheat spike of the cultivar BARI Gom 24 (Prodip) in Jhenaidah, Bangladesh, during 2016. The isolate was stored on dried filter paper at 4°C until experimentation (41). For reactivation, the strain was cultured on potato dextrose agar (PDA; 42 g/L) and grown at 25°C for 7–8 days. A small block of the MoT isolate BTJP 4 (5) was then transferred to fresh PDA medium and incubated under the same conditions (41). To stimulate sporulation, 10-day-old PDA cultures were rinsed under sterile conditions with 500 mL of deionized water to eliminate aerial hyphae, followed by incubation at 25–30°C for 2–3 days (8, 42). Conidia were harvested and suspended in sterile distilled water for microscopic observation. Germination was evaluated by monitoring conidial development under a compound microscope, and germinated conidia were manually counted and standardized to 1 × 10^5^ conidia/mL using a hemocytometer (Neubauer, 0.0025 mm²) (43). For the in vivo disease suppression assay, the susceptible wheat cultivar BARI Gom 26 was selected as the host plant.

### Chemicals and Preparation of Chemical Solution

3-methylpentanoic Acid was purchased from Sigma-Aldrich (product no. W343706). The fungicide Nativo® WG 75 (50:50 mixtures of trifloxystrobin and tebuconazole) was purchased in Dhaka, Bangladesh from Bayer Crop Science Ltd. A stock solution of 3-MP was prepared using a minimal amount of dimethyl sulfoxide (DMSO). Working concentrations of 25 µM, 50 µM, 75 µM, 100 µM, and 125 µM were subsequently formulated, ensuring that the final DMSO content did not exceed 1% (v/v), a level previously shown not to interfere with hyphal growth or sporulation of MoT (31).

### Bioassay with 3-MP on MoT Growth

3-MP was evaluated against MoT at different concentrations: 25 µM, 50 µM, 75 µM, 100 µM, and 125 µM. The commercial fungicide Nativo® 75 WG served as the industry standard. An autoclaved filter paper was affixed to the lid of each Petri dish, and the respective 3-MP concentration was applied to it. A 2 mm mycelial block of MoT was placed in the center of a PDA plate, and the lids were sealed tightly to prevent the loss of 3-MP. The radial mycelial growth (cm) of MoT was measured 7 days post-incubation. After this 7-day period, the Petri dish lids containing the 3-MP compound were replaced with new lids to eliminate its presence. The fungicidal effects of 3-MP were assessed by recording the radial growth (cm) of MoT up to 14 days post-incubation. Measurements were taken for the inhibition zones and fungal colony diameters influenced by the test compounds and the fungicide. Radial growth inhibition percentage (RGIP) (± standard error) (44) was calculated from mean values as:

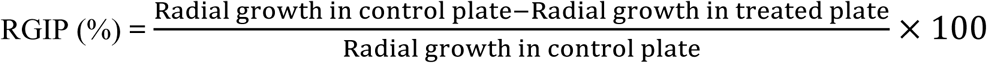

### Inhibition of Conidia Formation (Conidiogenesis)

The mycelium from a 10-day-old Petri dish culture of MoT was washed to deplete nutrients and promote conidiogenesis (8). The antifungal activity of 3-MP was evaluated at three concentrations (50 µM, 100 µM, and 150 µM) by applying the compound to autoclaved filter paper affixed to the Petri dish lid, which was then sealed tightly. A standard fungicidal suspension of Nativo® 75 WG was prepared in distilled water and used as a positive control. Negative control and treated Petri plates containing MoT were incubated at 28°C with >90% relative humidity (45). The light and dark periods were set to 14 hours and 10 hours, respectively. Conidiogenesis was examined after 48 hours using a Zeiss Primo Star microscope at 10X magnification, and images were captured with a Zeiss Axiocam ERc 5s. Each treatment was performed with three replication and the experiment was repeated three times.

### Inhibition of conidial germination and morphological alteration

The inhibitory effect of 3-MP on MoT conidial germination was assessed at three concentrations (25 µM, 50 µM, and 100 µM) over varying time intervals. A sterile filter paper moistened with sterile water was positioned at the base of a Petri dish to maintain humidity, alongside a microscope slide. A 20 µL droplet of MoT conidial suspension (1×10⁵ conidia/mL) was placed on the slide. Filter paper strips containing the specified 3-MP concentration were attached to the Petri dish lid’s inner surface, and the dish was securely sealed to minimize the loss of 3-MP. Incubation occurred at 25°C in darkness, with conidial germination monitored at 3, 6, 9, 12, and 24 h (46). For each treatment, 100 conidia from three replicates were analyzed using a Zeiss Primo Star microscope (40X magnification), with images recorded via a Zeiss Axiocam ERc 5s camera. The study included three replication and was repeated three times.

### Progression of wheat blast on detached wheat leaves

The detached leaf assay employed the wheat cultivar BARI Gom 26. Seeds were surface-sterilized using 3% sodium hypochlorite for 1 minute, rinsed thrice with sterilized distilled water and germinated on moistened filter paper in Petri dishes. Germinated seeds were transplanted into plastic pots containing a sand-compost-peat mixture (1:2:1 ratio). Plants were cultivated in a greenhouse under controlled conditions: 14/10 h light-dark cycle, 25°C (±2°C) temperature, and 65–70% relative humidity. At the five-leaf stage, leaves were excised and placed on 1% water agar supplemented with 10 mg/L kinetin (to delay senescence) in 90 mm Petri dishes (43). A 15 µL droplet of MoT conidial suspension (1×10⁶ conidia/mL) with 0.08% Tween 20 was applied to the adaxial (upper) surface of each leaf segment without wounding. Control leaves received sterile distilled water with Tween 20 (43). To evaluate the antifungal activity of 3-MP, three concentrations (25 µM, 50 µM, and 100 µM) were applied to autoclaved filter paper strips attached to the Petri dish lid, which was tightly sealed to prevent its loss. Plates were incubated at 28°C in darkness for 30 h, followed by continuous light exposure for 48 h. The experiment included three replicates per treatment and was repeated three times. Blast lesion diameters caused by MoT were measured on three leaves per treatment and concentration (47).

### Seedling infection assay (preventive and curative)

The seeds of wheat cultivar BARI Gom 26 were surface sterilized following the protocol described by Paul et al., (2022). Twenty seeds were sown in plastic pots (9 cm × 7 cm) containing sterilized field soil amended with NPK fertilizer. Fifteen days after seedling emergence, two separate experiments (preventive and curative assays) were conducted to evaluate seedling responses, with all pots arranged in a Completely Randomized Design for both experiments (35). For the preventive assay, autoclaved filter paper was glued to the inner upper side of an inverted plastic pot (250 mL), and 3-MP was applied at concentrations of 32 µM and 64 µM. The pots were sealed properly to ensure 3-MP retention. After 48 h of treatment, the 3-MP containing plastic pots were removed, and a conidial suspension of MoT 1×10^5^ conidia/mL was sprayed onto the seedlings until thoroughly wet. The plants were then covered with polythene bags to maintain high humidity (>90%) for 24 h at 25°C in darkness to promote fungal infection. For the curative assay, wheat seedlings were first spray-inoculated with MoT spores (1×10^5^ conidia/mL) and covered with an inverted plastic pot (250 mL). Autoclaved filter paper was glued to the upper side of the pot after 48 h post inoculation, and 3-MP was applied at concentrations of 16 µM, 32 µM, and 64 µM. The pots were sealed properly to retain 3-MP, and humidity was maintained at >90% for 24 hours at 25°C in darkness to facilitate fungal infection and after 24 h 3-MP was removed from the plastic cups. After treatment, all seedlings were transferred to a growth chamber set at 25°C with a 12 h light cycle and >85% relative humidity (31). Wheat plants that were untreated with MoT served as healthy controls, while plants inoculated with MoT but without 3-MP served as untreated controls. Sterilized water was sprayed daily on the inoculated seedlings at 4:00 PM to maintain high humidity. Disease development was assessed 7 days after inoculation, and each treatment was replicated three times. Disease severity was evaluated based on six progressive grades ranging from 0 to 5 (48). The scales were:

0= no lesions;

1= small brown, specks of pinhead size;

2= small, roundish to slightly elongated necrotic, gray spots about 1–2mm in diameter; 3= typical blast lesions infecting 51% of leaf area and many dead leaves.

4= large lesion

5= complete shriveling of leaf blades

The intensity of the disease (DI) was determined by employing the formula:

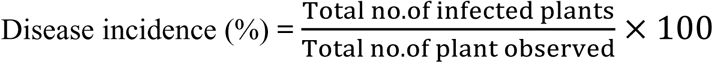

After that, the disease severity was calculated using the following formula:

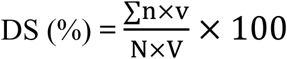

Where,

DS= Disease Severity.

n= Number of leaf infected.

v= Value score of each category attack.

N= Number of leaves observed

V= Value of highest score

### Spike Infection Assay

The seeds of the wheat BARI Gom 26 were surface sterilized by following the protocol described by Paul et al. (2022). Twenty seeds were sown in plastic pots (20 cm x 18 cm) containing sterilized field soil amended with NPK fertilizer and grown for 61 days in a greenhouse under 12/12-hour light–dark cycles with 65% relative humidity. All pots were arranged in a Completely Randomized Design for this experiment. Wheat spikes were inoculated with MoT spores (1 x 10⁵ conidia/mL) and covered with plastic bags to ensure infection. Five autoclaved filter papers were glued onto the upper and lateral sides of each plastic bag, and 3-MP was applied at concentrations of 8 µM and 16 µM. The bags were sealed properly to prevent 3-MP loss, and humidity (>90%) was maintained for 24 h at 25°C in darkness to facilitate fungal infection. After 24 h, all 3-MP were removed from the plastic pots ang bags. After that, the plants were transferred to a growth chamber set at 25°C with a 12 h light cycle and >85% relative humidity. Wheat plants treated without MoT served as healthy controls, while plants treated with MoT but without 3-MP served as untreated controls. To maintain high humidity, sterilized water was sprayed on the inoculated plants daily at 4:00 PM. Disease development was assessed 10 days after inoculation, and each treatment was replicated three times.

### MoT mycelial viability test using fluorescein diacetate (FDA)

The viability of fungal cells was assessed using FDA staining. At the untreated control, live fungal cells with intact membranes exhibit green fluorescence (49). Hyphae of MoT treated with 3-MP at concentrations ranging from 50 µM to 125 µM, were carefully collected from culture plates. The hyphae were resuspended in 10 mM phosphate-buffered saline (PBS, pH 7.4) containing 100 μg/mL FDA. Samples were incubated in the dark at 25°C for 10 minutes to allow FDA hydrolysis by viable cells. After incubation, the hyphae were washed twice with 1 mL of 1x PBS (pH 7.4) to remove excess stain (49). Stained samples were observed under a Zeiss Primo Star microscope to detect green fluorescence indicating cell viability. FDA-derived fluorescein was excited using a 488 nm argon laser, and emission was collected at approximately 520 nm (green channel). Each field was first focused under bright-field to locate the mycelium before switching to fluorescence mode for imaging. And images were captured using a Zeiss Axiocam ERc 5s camera.

### Essential MoT protein identification and interaction with 3-MP

To identify the key proteins within the MoT, we conducted a Basic Local Alignment Search Tool for proteins (BLASTp) analysis, comparing the MoT proteome against a database of essential proteins. Our BLASTp analysis were performed with an e-value threshold set at 10^−300^, a sequence identity of 35%, a bit score of 100, and others standard parameter (50). Subsequently, we investigated the interactions between 3-MP and the identified essential proteins using vina.bat scripts executed via PowerShell on a Linux platform (51). We selected four promising complexes for 200 ns molecular dynamics simulations using Desmond package in Schrodinger suite v2023-4 on a high-performance computing system with a Supermicro cluster CPU and NVIDIA RTX GPU. Before simulating the protein-ligand complexes protein preparation steps were incorporated with preprocessing refinement and geometry optimization. Subsequently, the system builder was utilized to prepare the protein-ligand complex. The SPC water model was chosen, configured in an orthorhombic shape after volume minimization, with 10 Å × 10 Å × 10 Å periodic boundary conditions applied along the x, y, and z axes of the protein-ligand complex. Additionally, to neutralize the system Na^+^ ions were added with excluding ion and salt placement within 20 Å to maintain this neutrality. In the next step the OPLS2005 force field was utilized to minimize the complex’s energy through heating and equilibrium processes before running the MD simulations. Running on an NVIDIA RTX GPU, each 200 ns simulation took approximately 6.5 hours numerical values were collected and analyzed using a simulation interaction diagram to study trajectories and fluctuations.

### RNA extraction and reverse transcription-quantitative polymerase chain reaction (RT-qPCR)

A 30 µL droplet of Potato Dextrose Broth (PDB) containing MoT mycelia was placed on a sterile glass slide inside a Petri plate and incubated at 25 °C for 3 days. Following incubation, mycelia were treated with two concentrations of 3-MP, 100 µM and 150 µM. Samples were collected at 0, 24, and 48 h post–treatment and immediately frozen in liquid nitrogen for subsequent RNA extraction. Total RNA was isolated from MoT mycelia using Trizol reagent (Invitrogen, USA). RNA concentration and purity were assessed using a NanoDrop spectrophotometer (NanoDrop Technologies, USA). First-strand cDNA was synthesized from up to 5 µg of RNA using the GoScript™ Reverse Transcription System (Promega, USA) following the manufacturer’s instructions. Briefly, RNA–primer mixtures were denatured at 70 °C for 5 min, chilled on ice, and combined with reverse transcription reagents to a final volume of 20 µL. Reactions were incubated at 25 °C for 5 min, extended at 42 °C for 60 min, and terminated by heating at 70 °C for 15 min. Quantitative PCR (qPCR) was carried out using GoTaq® qPCR Master Mix (Promega, USA) with gene-specific primers (**Table 4**), employing actin (MGG_03982) as the internal reference on an Analytik Jena qTower 3 84G PCR Thermal Cycler. Each 20 µL reaction contained 10 µL master mix, 1 µL forward primer, 1 µL reverse primer, 1 µL cDNA template, and 7 µL nuclease-free water. The cycling program consisted of an initial denaturation at 95 °C for 3 min, followed by 45 cycles of 95 °C for 15 s, 58 °C for 15 s, and 60 °C for 20 s, with a final hold at 4 °C. Relative gene expression levels were calculated using the 2^−ΔΔCT^ method (52), and results were expressed as log₂ fold change.

**Table 4:**
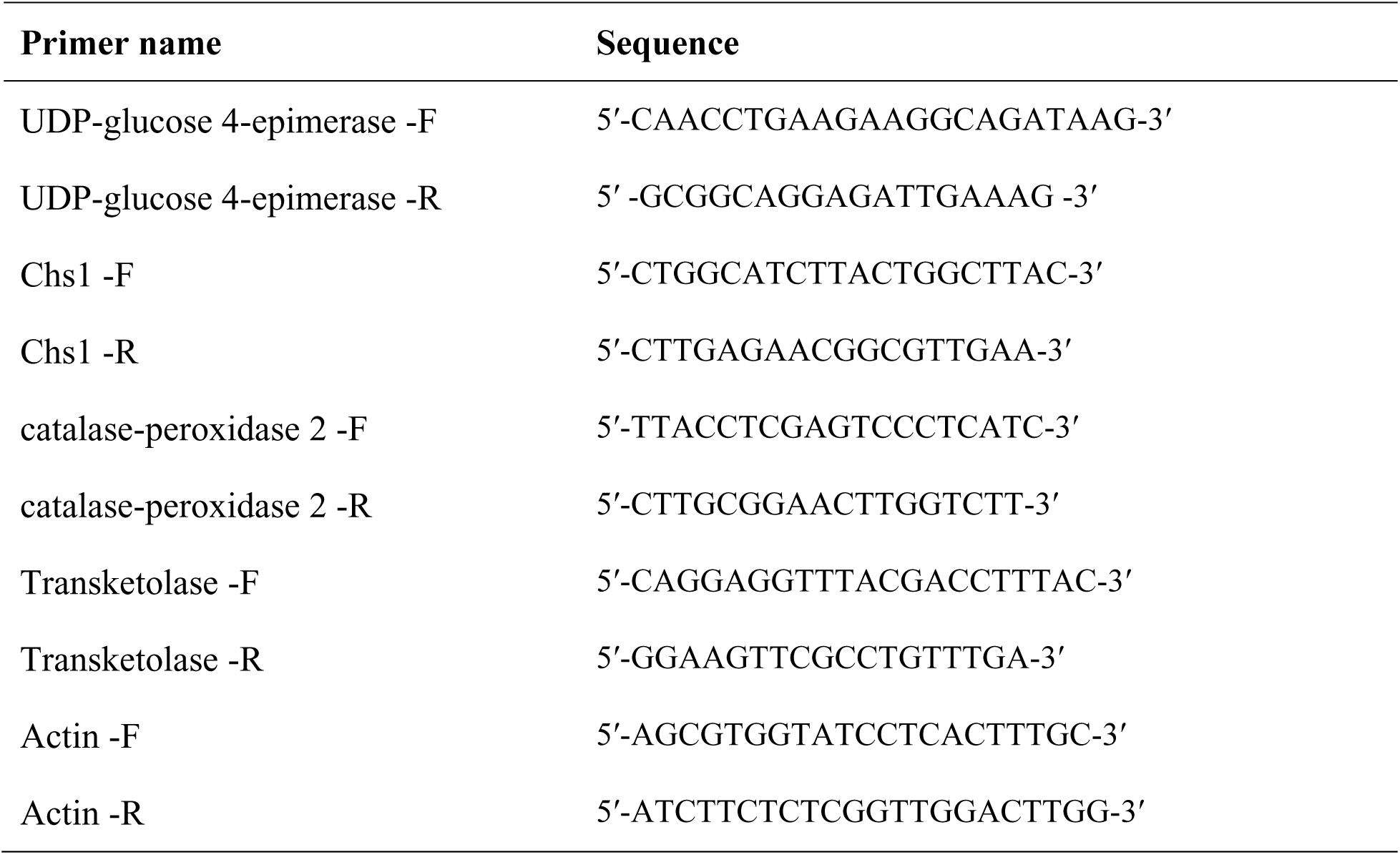
List of genes and gene-specific primers used in the RT-qPCR analysis

## Statistical analysis

Experiments were arranged in a completely randomized design to evaluate the biological activity of 3-MP in comparison to a standard fungicide. Data were analyzed using one-way analysis of variance (ANOVA), and treatment means were compared using Tukey’s honest significant difference (HSD) post hoc test at a significance level of p ≤ 0.01. All statistical analyses were performed using R software (version 4.5.1, accessed 5 December, 2025) integrated into RStudio (version 2025.09.1+401, accessed 5 December, 2025). Results are presented as mean ± standard error (SE) from 3 biological replicates in tables and figures.

## Data availability

All data are included in the article or in the supporting information.

## Supporting information

This article contains supporting information. Figure S1, Figure S2.

## Acknowledgments

The authors are thankful to Bangladesh Academy of Sciences for funding this work through a project under BAS-USDA-PAL funding system to T.I.

## Author contributions

M.S.P.S writing–original draft; methodology; validation; investigation; Digital PCR gene expression analysis; visualization; software; formal analysis; data curation; S.H software; molecular dynamic simulation analysis; Writing – review and editing; R.B.A Methodology, Validation, Writing – review and editing; D.R.G Supervision, Resources, Validation, Writing – review and editing; M.T.I, Conceptualization, Funding acquisition, Project administration, Resources, Supervision, Validation, Writing – review and editing.

## Funding and additional information

This research was supported by funding partially from the Bangladesh Academy of Sciences, Bangladesh under a BAS-USDA-PAL project to T.I. and the University Grants Commission of Bangladesh under the project number Agriculture (Crop Science-35/2022-23) to D.R.G.

## References

1. Savary, S., Willocquet, L., Pethybridge, S. J., Esker, P., McRoberts, N., and Nelson, A. (2019) The global burden of pathogens and pests on major food crops. *Nat*. Ecol. Evol. 3, 430–439

2. Islam, M. T., Croll, D., Gladieux, P., Soanes, D. M., Persoons, A., Bhattacharjee, P., Hossain, M. S., Gupta, D. R., Rahman, M. M., Mahboob, M. G., Cook, N., Salam, M. U., Surovy, M. Z., Sancho, V. B., Maciel, J. L. N., Nhani Júnior, A., Castroagudín, V. L., Reges, J. T. de A., Ceresini, P. C., Ravel, S., Kellner, R., Fournier, E., Tharreau, D., Lebrun, M. H., McDonald, B. A., Stitt, T., Swan, D., Talbot, N. J., Saunders, D. G. O., Win, J., and Kamoun, S. (2016) Emergence of wheat blast in Bangladesh was caused by a South American lineage of *Magnaporthe oryzae*. BMC Biol. 14, 84

3. Tembo, B., Mulenga, R. M., Sichilima, S., M’siska, K. K., Mwale, M., Chikoti, P. C., Singh, P. K., He, X., Pedley, K. F., Peterson, G. L., Singh, R. P., and Braun, H. J. (2020) Detection and characterization of fungus (*Magnaporthe oryzae* pathotype *Triticum*) causing wheat blast disease on rain-fed grown wheat (*Triticum aestivum* L.) in Zambia. PLoS One. 10.1371/journal.pone.0238724

4. Kamoun, S., Talbot, N. J., and Islam, M. T. (2019) Plant health emergencies demand open science: Tackling a cereal killer on the run. PLoS Biol. 17, e3000302

5. Islam, M. T., Kim, K. H., and Choi, J. (2019) Wheat blast in Bangladesh: The current situation and future impacts. Plant Pathol. J. 35, 1–10

6. Pequeno, D. N. L., Ferreira, T. B., Fernandes, J. M. C., Singh, P. K., Pavan, W., Sonder, K., Robertson, R., Krupnik, T. J., Erenstein, O., and Asseng, S. (2024) Production vulnerability to wheat blast disease under climate change. *Nat*. Clim. Change 14, 178–183

7. Wilson, R. A., and Talbot, N. J. (2009) Under pressure: investigating the biology of plant infection by *Magnaporthe oryzae*. Nat. Rev. Microbiol. 7, 185–195

8. Urashima, A. S., Igarashi, S., and Kato, H. (1993) Host range, mating type, and fertility of *Pyricularis grisea* from wheat in Brazil. Plant Dis. 77, 1211–1216

9. Vicentini, S. N. C., Casado, P. S., de Carvalho, G., Moreira, S. I., Dorigan, A. F., Silva, T. C., Silva, A. G., Custódio, A. A. P., Gomes, A. C. S., Maciel, J. L. N., Hawkins, N., McDonald, B. A., Fraaije, B. A., and Ceresini, P. C. (2022) Monitoring of Brazilian wheat blast field populations reveals resistance to QoI, DMI, and SDHI fungicides. Plant Pathol. 71, 304–321

10. Mohiddin, F. A., Bhat, N. A., Wani, S. H., Bhat, A. H., Ahanger, M. A., Shikari, A. B., Sofi, N. R., Parveen, S., Khan, G. H., Bashir, Z., Vachova, P., Hassan, S., and Sabagh, A. E. L. (2021) Combination of strobilurin and triazole chemicals for the management of blast disease in mushk budji-aromatic rice. J. Fungi. 10.3390/jof7121060

11. Kohli, M. M., Cazal, C., and Chavez, A. (2020) Integrated management of wheat blast disease. In *Wheat Blast*. 10.1201/9780429470554-10

12. Castroagudín, V. L., Ceresini, P. C., de Oliveira, S. C., Reges, J. T. A., Maciel, J. L. N., Bonato, A. L. V., Dorigan, A. F., and McDonald, B. A. (2015) Resistance to QoI fungicides is widespread in Brazilian populations of the wheat blast pathogen *Magnaporthe oryzae*. Phytopathology 105, 284–294.

13. Yoshioka, M., Shibata, M., Morita, K., Islam, M. T., Fujita, M., Hatta, K., Tougou, M., Tosa, Y., and Asuke, S. (2024) Breeding of a near-isogenic wheat line resistant to wheat blast at both seedling and heading stages through incorporation of Rmg8. Phytopathology 114, 1843–1850

14. Singh, P. K., Gahtyari, N. C., Roy, C., Roy, K. K., He, X., Tembo, B., Xu, K., Juliana, P., Sonder, K., Kabir, M. R., and Chawade, A. (2021) Wheat blast: A disease spreading by intercontinental jumps and its management strategies. Front. Plant Sci. 12, 710707

15. Zhang, R. X., Zhang, Y. F., Yang, H., Zhang, X. D., Yang, Z. G., Li, B. B., Sun, W. H., Yang, Z., Liu, W. T., and Chen, K. M. (2025) An optimized editing approach for wheat genes by improving sgRNA design and transformation strategies. Int. J. Mol. Sci. 10.3390/ijms26083796/S1

16. Kwon, Y. S., Ryu, C. M., Lee, S., Park, H. B., Han, K. S., Lee, J. H., Lee, K., Chung, W. S., Jeong, M. J., Kim, H. K., and Bae, D. W. (2010) Proteome analysis of *Arabidopsis* seedlings exposed to bacterial volatiles. Planta 232, 1355–1370

17. Morath, S. U., Hung, R., and Bennett, J. W. (2012) Fungal volatile organic compounds: A review with emphasis on their biotechnological potential. Fungal Biol. Rev. 26, 73–83

18. Mercier, J., and Smilanick, J. L. (2005) Control of green mold and sour rot of stored lemon by biofumigation with *Muscodor albus*. Biol. Control 32, 401–407

19. Zuo, G., Han, Y., Dong, Q., Lu, W., Gao, C., Zhao, N., Liu, S., Fan, G., Liu, C., and Xiang, W. (2025) Antifungal activity and mechanism of volatile organic compounds produced by *Bacillus siamensis* NEAU-ZGX24 against *Botrytis cinerea*. Microbiol. Res. 299, 128265

20. Yang, Y., Ma, W., Johnson, E. T., Xie, Y., and Zhao, R. (2025) Antifungal effects of three natural branched medium-chain fatty acids and their potential as fumigants against *Aspergillus flavus* in stored peanut seeds. Food Control 168, 110950

21. . Castroagudín, V. L., Ceresini, P. C., de Oliveira, S. C., Reges, J. T., Maciel, J. L., Bonato, A. L., … & McDonald, B. A. (2015). Resistance to QoI fungicides is widespread in Brazilian populations of the wheat blast pathogen *Magnaporthe oryzae*. Phytopathology, 105(3), 284–294.

22. Gupta, D. R., Paul, S. K., Ino, M., Oowatari, Y., and Ueno, M. (2025) *Bacillus safensis* NI2B displays strong antagonistic activities against rice blast pathogen *Pyricularia oryzae* through the production of volatile compounds. Phytopathol. Res. 10.1186/s42483-025-00340-6

23. Paul, S. K., Gupta, D. R., Ino, M., Sujon, M. S. P., and Ueno, M. (2025) 3-Methyl pentanoic acid suppresses gray mold disease potentially targeting cell-wall integrity (CWI) and mitogen-activated protein kinase (MAPK) pathways in *Botrytis cinerea*. BMC Microbiol. 25, 1–11

24. Wijaya, C. H., Ulrich, D., Lestari, R., Schippel, K., and Ebert, G. (2005) Identification of potent odorants in different cultivars of snake fruit (*Salacca zalacca* (Gaert.) Voss) using gas chromatography-olfactometry. J. Agric. Food Chem. 53, 1637–1641

25. Cohen, S. M., Eisenbrand, G., Fukushima, S., Gooderham, N. J., Guengerich, F. P., Hecht, S. S., Rietjens, I. M. C. M., Rosol, T. J., Harman, C., and Taylor, S. V. (2020) GRAS 29 flavoring substances. Food Technol. 74, 44–51

26. Singh, V., Satheesh, S. V., Raghavendra, M. L., and Sadhale, P. P. (2007) The key enzyme in galactose metabolism, UDP-galactose-4-epimerase, affects cell-wall integrity and morphology in Candida albicans even in the absence of galactose. Fungal Genet. Biol. 44, 563–574

27. Park, J., Tefsen, B., Arentshorst, M., Lagendijk, E., van den Hondel, C. A., van Die, I., and Ram, A. F. (2014) Identification of the UDP-glucose-4-epimerase required for galactofuranose biosynthesis and galactose metabolism in *Aspergillus niger*. Fungal Biol. Biotechnol. 1, 6

28. Fernandez, J., Marroquin-Guzman, M., and Wilson, R. A. (2014) Evidence for a transketolase-mediated metabolic checkpoint governing biotrophic growth in rice cells by the blast fungus *Magnaporthe oryzae*. PLoS Pathog. 10.1371/journal.ppat.1004354

29. Wang, D., Peng, C., Zheng, X., Chang, L., Xu, B., and Tong, Z. (2020) Secretome analysis of the banana *Fusarium* wilt fungi Foc R1 and Foc TR4 reveals a new effector OASTL required for full pathogenicity of Foc TR4 in banana. Biomolecules 10, 1–17

30. Surovy, M. Z., Rahman, S., Rostás, M., Islam, T., and von Tiedemann, A. (2023) Suppressive effects of volatile compounds from *Bacillus* spp. on *Magnaporthe oryzae Triticum* (MoT) pathotype, causal agent of wheat blast. Microorganisms 11, 1291. doi: 10.3390/microorganisms11051291.

31. Paul, S. K., Chakraborty, M., Rahman, M., Gupta, D. R., Mahmud, N. U., Rahat, A. A. M., Sarker, A., Hannan, M. A., Rahman, M. M., Akanda, A. M., Ahmed, J. U., and Islam, T. (2022) Marine natural product antimycin A suppresses wheat blast disease caused by *Magnaporthe oryzae* Triticum. J. Fungi. 10.3390/jof8060618

32. Tyagi, S., Lee, K. J., Shukla, P., and Chae, J. C. (2020) Dimethyl disulfide exerts antifungal activity against *Sclerotinia minor* by damaging its membrane and induces systemic resistance in host plants. Sci. Rep. 10, 6547

33. Tilocca, B., Cao, A., and Migheli, Q. (2020) Scent of a killer: Microbial volatilome and its role in the biological control of plant pathogens. Front. Microbiol. 11, 41

34. Chevalier, L., Pinar, M., Le Borgne, R., Durieu, C., Peñalva, M. A., Boudaoud, A., and Minc, N. (2023) Cell wall dynamics stabilize tip growth in a filamentous fungus. PLoS Biol. 21, e3001981

35. Chakraborty, M., Mahmud, N. U., Muzahid, A. N. M., Rabby, S. M. F., and Islam, T. (2020) Oligomycins inhibit *Magnaporthe oryzae Triticum* and suppress wheat blast disease. PLoS One. 10.1371/journal.pone.0233665

36. Lee, M. J., Gravelat, F. N., Cerone, R. P., Baptista, S. D., Campoli, P. V., Choe, S.-I., Kravtsov, I., Vinogradov, E., Creuzenet, C., Liu, H., Berghuis, A. M., Latgé, J.-P., Filler, S. G., Fontaine, T., and Sheppard, D. C. (2013) Overlapping and distinct roles of *Aspergillus fumigatus* UDP-glucose 4-epimerases in galactose metabolism and the synthesis of galactose-containing cell wall polysaccharides. J. Biol. Chem. 289, 1243–1256

37. Thoden, J. B., Wohlers, T. M., Fridovich-Keil, J. L., and Holden, H. M. (2001) Human UDP-galactose 4-epimerase. J. Biol. Chem. 276, 15131–15136

38. El-Ganiny, A. M., Sheoran, I., Sanders, D. A. R., and Kaminskyj, S. G. W. (2010) *Aspergillus nidulans* UDP-glucose-4-epimerase UgeA has multiple roles in wall architecture, hyphal morphogenesis, and asexual development. Fungal Genet. Biol. 47, 629–635

39. Qu, Z., Chen, D., Hu, H., Liu, H., Zheng, L., Huang, J., Li, Y., Zhu, L., and Chen, X. (2026) Selectively targeting UDP-glucose 4-epimerase *MoUGE1* for controlling rice blast disease. J. Adv. Res. In press. doi: 10.1016/j.jare.2026.01.072.

40. Montejano-Ramírez, V., Ávila-Oviedo, J. L., Campos-Mendoza, F. J., and Valencia-Cantero, E. (2024) Microbial volatile organic compounds: Insights into plant defense. Plants 13, 2013. 10.3390/plants13152013

41. Gupta, D. R., Surovy, M. Z., Mahmud, N. U., Chakraborty, M., Paul, S. K., Hossain, M. S., Bhattacharjee, P., Mehebub, M. S., Rani, K., Yeasmin, R., Rahman, M., and Islam, M. T. (2020) Suitable methods for isolation, culture, storage, and identification of wheat blast fungus *Magnaporthe oryzae* Triticum pathotype. Phytopathol. Res. 10.1186/s42483-020-00070-x

42. Urashima, A. S., Hashimoto, Y., Don, L. D., Kusaba, M., Tosa, Y., Nakayashiki, H., and Mayama, S. (1999) Molecular analysis of the wheat blast (*Pyricularia oryzae*) population in Brazil with a homolog of retrotransposon MGR583. Ann. Phytopathol. Soc. Jpn. 65, 429–436

43. Kumar, K., Xi, K., Turkington, T. K., Tekauz, A., Helm, J. H., and Tewari, J. P. (2011) Evaluation of a detached leaf assay to measure fusarium head blight resistance components in barley. Can. J. Plant Pathol. 33, 364–374

44. Riungu, G. M., Muthomi, J. W., Narla, R. D., Wagacha, J. M., and Gathumbi, J. K. (2008) Management of fusarium head blight of wheat and deoxynivalenol accumulation using antagonistic microorganisms. Plant Pathol. J. (Faisalabad) 7, 13–19

45. Rabby, S. M. F., Chakraborty, M., Gupta, D. R., Rahman, M., Paul, S. K., Mahmud, N. U., Rahat, A. A. M., Jankuloski, L., and Islam, T. (2022) Bonactin and feigrisolide C inhibit *Magnaporthe oryzae* Triticum fungus and control wheat blast disease. Plants. 10.3390/plants11162108

46. Chakraborty, M., Rabby, S. M. F., Gupta, D. R., Rahman, M., Paul, S. K., Mahmud, N. U., Rahat, A. A. M., Jankuloski, L., and Islam, T. (2022) Natural protein kinase inhibitors, staurosporine, and chelerythrine suppress wheat blast disease caused by *Magnaporthe oryzae* Triticum. Microorganisms. 10.3390/microorganisms10061186

47. Ha, X., Koopmann, B., and von Tiedemann, A. (2016) Wheat blast and fusarium head blight display contrasting interaction patterns on ears of wheat genotypes differing in resistance. Phytopathology 106, 270–281

48. Hyon, G. S., Nga, N. T. T., Chuma, I., Inoue, Y., Asano, H., Murata, N., Kusaba, M., and Tosa, Y. (2012) Characterization of interactions between barley and various host-specific subgroups of *Magnaporthe oryzae* and *M. grisea*. J. Gen. Plant Pathol. 78, 237–246

49. Jones, K., Kim, D. W., Park, J. S., and Khang, C. H. (2016) Live-cell fluorescence imaging to investigate the dynamics of plant cell death during infection by the rice blast fungus *Magnaporthe oryzae*. BMC Plant Biol. 16, 1–8

50. Hasnat, S., Hoque, M. N., Mahbub, M. M., Sakif, T. I., Shahinuzzaman, A. D. A., and Islam, T. (2024) Pantothenate kinase: A promising therapeutic target against pathogenic *Clostridium* species. Heliyon 10, e34544

51. Hasnat, S., Rahman, S., Alam, M. B., Suin, F. M., Yeasmin, F., Suha, T., Supty, N. T., Sabila, S., Chowdhury, A., Shahinuzzaman, A. D. A., Mahbub, M. M., Islam, T., and Hoque, M. N. (2025) High throughput screening identifies potential inhibitors targeting trimethoprim-resistant DfrA1 protein in *Klebsiella pneumoniae* and *Escherichia coli*. Sci. Rep. 15, 7141

52. Livak, K. J., and Schmittgen, T. D. (2001) Analysis of relative gene expression data using real-time quantitative PCR and the 2^−ΔΔCT^ method. Methods 25, 402–408

